# Absolute quantification of transcription factors reveals principles of gene regulation in erythropoiesis

**DOI:** 10.1101/812123

**Authors:** Mark A. Gillespie, Carmen G. Palii, Daniel Sanchez-Taltavull, Paul Shannon, William J.R. Longabaugh, Damien J. Downes, Karthi Sivaraman, Jim R. Hughes, Nathan D. Price, Theodore J. Perkins, Jeffrey A. Ranish, Marjorie Brand

## Abstract

Dynamic cellular processes such as differentiation are driven by changes in the abundances of transcription factors (TFs). Yet, despite years of studies we still do not know the protein copy number of TFs in the nucleus. Here, by determining the absolute abundances of 103 TFs and co-factors during the course of human erythropoiesis, we provide a dynamic and quantitative scale for TFs in the nucleus. Furthermore, we establish the first Gene Regulatory Network of cell fate commitment that integrates temporal protein stoichiometry data with mRNA measurements. The model revealed quantitative imbalances in TFs cross-antagonistic relationships that underlie lineage determination. Finally, we made the surprising discovery that in the nucleus, corepressors are dramatically more abundant than coactivators at the protein, but not at the RNA level, with profound implications for understanding transcriptional regulation. These analyses provide a unique quantitative framework to understand transcriptional regulation of cell differentiation in a dynamic context.

## Introduction

Quantitative changes in TF abundances drive dynamic cellular processes such as differentiation by activating lineage-specific gene expression programs and simultaneously repressing competing lineages (Graf and Enver, 2009; Orkin and Zon, 2008). At the mechanistic level, biochemical studies have shown that TFs function through the recruitment and/or stabilization of cofactors such as chromatin modifiers to target genes (Brand et al., 2019; Demers et al., 2007). On a more global scale, genomic and transcriptomic studies have uncovered intricate gene regulatory relationships whereby specific combinations of TFs cooperate or compete to regulate cell-specific gene programs (Reiter et al., 2017). The complexity of these relationships is best captured using gene regulatory networks (GRNs) to model the activating and repressing roles of TFs, which underlie lineage fate decisions in multipotent cells such as hematopoietic progenitors (Gottgens, 2015; Novershtern et al., 2011; Rothenberg, 2019; Swiers et al., 2006).

In contrast to the current profusion of transcriptomic data, large-scale quantitative proteomic information is scarce for low abundance proteins such as TFs and cofactors, particularly in human stem/progenitor cells. This critical lack of quantitative proteomic data is a major impediment to addressing fundamental questions in transcriptional regulation, such as TF and cofactor availability in the nucleus (Schmidt et al., 2016). Lack of quantitative proteomic data is also problematic for understanding dynamic processes such a cell fate decisions or differentiation that are based on changes in the stoichiometry of lineage-specifying (LS)-TFs (Graf and Enver, 2009; Orkin and Zon, 2008; Palii et al., 2019). In that regard it is particularly striking that none of the current GRNs for hematopoiesis incorporate quantitative protein abundance data for TFs.

Integration of proteomic data into GRNs is a significant challenge as it requires the ability to measure the abundances of multiple proteins relative to one another (i.e. stoichiometry) within the same sample (Vitrinel et al., 2019), information that cannot easily be obtained with antibody-based methods. Moreover, for dynamic GRNs, the same proteins must be repeatedly measured over time in different samples. While mass spectrometry (MS) is a powerful method for protein identification, absolute quantification is more challenging. A number of approaches have been described for absolute quantification, including the use of synthetic isotopically labeled (SIL) peptides as internal standards, which are typically used for quantification of relatively small numbers of proteins, and label-free methods for large scale quantification [see (Liu et al., 2016) for review]. Both stable isotope dilution (SID) and label-free approaches can be used in conjunction with targeted MS approaches, such as selected reaction monitoring (SRM) which are well-suited for reproducible and quantitative measurements of a defined set of analytes over a wide dynamic range of abundances. However, thus far no one has developed and deployed protein detection methods with the required sensitivity and reproducibility to systematically determine the absolute abundances of endogenous TFs and cofactors in rare stem and progenitor cells.

Here, we developed targeted SRM assays in hematopoietic and erythroid nuclear extracts and applied these assays along with SIL peptides as internal standards. This allowed us to systematically determine the absolute abundances of 103 endogenous TFs and co-factors at 13 sequential time points as human hematopoietic stem and progenitor cells (HSPCs) differentiate along the pathway to erythroid cells. In addition to defining the range of protein concentration (copy number per nucleus) for master regulators of hematopoiesis and erythropoiesis, the data revealed surprising differences in stoichiometry and dynamics between TFs, coactivators and corepressors that occur at the protein level, but not at the RNA level with important implications for understanding gene regulation during differentiation. Finally, through mathematical modeling we generated for the first time a dynamic regulatory network of erythroid commitment from hematopoietic stem cells (HSCs) that integrates quantitative changes in RNA and protein levels (stoichiometry) of TFs over time.

## Results

### Absolute Quantification of Transcription Factors during Human Erythropoiesis

While proteomic studies have been performed in erythroid cells (Amon et al., 2019; Brand et al., 2004; Gautier et al., 2016; Jassinskaja et al., 2017; Liu et al., 2017), no previous study has used targeted MS approaches to provide systematic and absolute quantification of TF proteins with simultaneous mRNA measurements during the dynamic process of erythroid differentiation. Furthermore, previous MS-based studies have not measured TFs prior to lineage commitment, preventing the study of quantitative changes in TFs that underlie cell fate decisions. To address this, we employed a well-characterized ex vivo culture system in which cord blood-derived human multipotent HSPCs are induced to differentiate along the erythroid lineage (Giarratana et al., 2005; Palii et al., 2011a). Previously, using single cell mass cytometry, we showed that in this system, cells recapitulate all sequential stages of erythropoiesis over time, including multipotent progenitors (MPP) at day 0, common myeloid progenitors (CMP) and megakaryocyte-erythroid progenitors (MEP) at days 2-4, erythroid progenitors “colony-forming-unit erythroid” (CFU-E) at days 6-11, followed by terminally differentiating precursors: pro-erythroblasts (ProEB) at day 12, basophilic erythroblasts (Baso_EB) at day 14 and polychromatophilic erythroblasts (Poly_EB) at day 16 (Palii et al., 2019). Thus, this provides an ideal system for quantitative proteomic and transcriptomic analyses. Cells were harvested at 13 sequential time points and processed in parallel for RNA sequencing (RNAseq) and nuclear protein extraction (Figure S1A). First, RNAseq analysis revealed a main trajectory along two principal components from day 0 to day 16 (Figure S1C). Furthermore, a correlation heatmap indicated that the transcriptome changes gradually throughout the time series with sharper changes at day 2 (transition to CMP/MEP), day 11.5 (transition to ProEB) and day 14 (transition to Baso_EB) (Figures 1E and S2A). For proteomic analyses, nuclear extracts were first analyzed using an unbiased data-dependent MS approach with isobaric 8-plex iTRAQ reagents (Ross et al., 2004) which led to the identification and relative quantification of 3,905 proteins over time (<u>Tables S1</u> and <u>S2</u>, Figure S1E). Analysis of iTRAQ data allowed identification and relative quantification of 3,905 proteins, including 655 TFs, over time (<u>Table S1</u>). K-means clustering with gene ontology (GO) analyses revealed groups of proteins with enrichment in erythroid-related categories (e.g. “O_2_ transport” in Baso_EB (cluster 8), “cell cycle” in the highly proliferative CFU-E populations (cluster 6)) (Figure S1E and Table S2). Furthermore, the GO term “eukaryotic translation initiation” was found enriched in proteins that increase at the ProEB stage (day 12), consistent with the importance of protein translation in terminal erythroid differentiation (Alvarez-Dominguez et al., 2017). Interestingly, we found that proteins involved in oxidative phosphorylation (cluster 3) increase at day 2, which coincides with MPPs commitment to the myeloid/erythroid lineage. Given that oxidative phosphorylation correlates with lineage commitment (Oburoglu et al., 2016), this finding further supports the relevance of our ex vivo differentiation system to reveal molecular events that occur in early hematopoietic progenitors.

**Figure 1.**
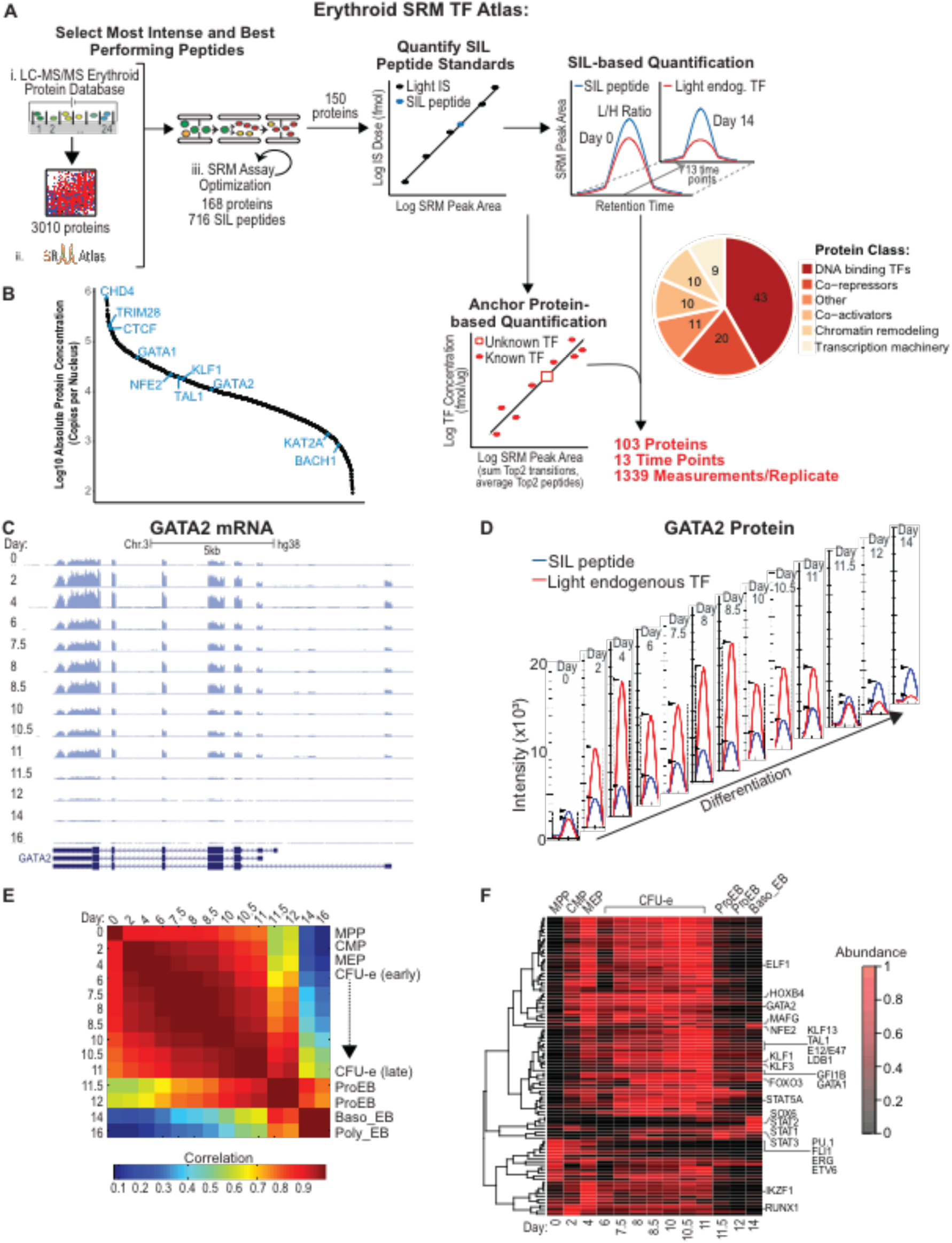
Absolute Quantification of Transcription Factors and Co-factors during Human Erythropoiesis. (A) Schematic of SRM assay development to quantify TFs and cofactors during hemato/erythropoiesis. Absolute abundance was determined at each time-point using either the SIL-based or the Anchor protein–based quantification method. Two biological replicates were performed. SIL, stable isotope labeled; IS, internal standard. (B) Range of protein abundances in the nucleus averaged for all time points. (C) RNA-seq coverage of the GATA2 gene over time as displayed by the UCSC genome browser. Replicate 1 is shown as a representative example. (D) Quantification of the GATA2 protein over time by SRM. Ion chromatograms of endogenous (red) and SIL (blue) EVSPDPSTTGAASPASSSAGGSAAR peptide from the Skyline software. Replicate 1 is shown as a representative example. (E) Correlation matrix between RNA-seq experiments at the indicated days. The heatmap displays Pearson correlations. The stages of differentiation (indicated on the right) were attributed to the time-points using CyTOF as previously described (Palii et al., 2019). (F) k-means clustering analysis of normalized protein abundances at the indicated days. Protein abundances were measured by SRM and are shown as a heatmap after normalization. See also Figures S1, S2, S3 and Tables S1, S2, S3 and S4.

To systematically determine the absolute concentration of TFs and cofactors, we employed SRM-based targeted MS together with SIL peptide-based quantification (Figure 1A). First, we established an Erythroid SRM TF Atlas consisting of parameters needed to monitor 411 isotopically heavy and light peptide pairs, corresponding to 150 proteins (1-4 peptide pairs per protein) (<u>Tables S3</u>, <u>S4</u> and <u>Methods</u>). SRM assay development, including selection of proteotypic peptides, transition ions, and the linear ranges of quantification is described in Methods. Absolute quantification was then achieved by a combination of SID and label free quantification for 103 proteins that span various categories (e.g. DNA binding TFs, co-activators, co-repressors, chromatin modifiers) at 13 sequential time-points from MPP to basophilic erythroblasts (Figure 1A, S2B, S3; Table S4; see Methods). Comparing SRM results with iTRAQ measurements, we found a good correlation in protein changes during differentiation (Figure S2C). Importantly, SRM measurements covered a wide dynamic range of nuclear protein abundances, ranging from less than 500 copies for some factors (e.g. BACH1, GATA2, KAT2A) to above 100,000 copies for others (e.g. CTCF, TRIM28/KAP1, CHD4) (Figure 1B).

Using clustering analysis, we identified several groups of temporally-regulated proteins (Figure 1F). Interestingly, all master regulators of erythropoiesis (GATA1, TAL1, KLF1, KLF3, GFI1B, STAT5A) are characterized by a gradual increase from MPP to late CFU-E, followed by a sharp decline starting at the ProEB stage that marks the beginning of terminal erythroid differentiation (Figure 1D, F). This decrease was confirmed by Western blot (Figure S1D). While most factors are present at low levels in terminally differentiated erythroid cells, some TFs are more abundant, including NFE2, MAFG, FOXO3, SOX6, STAT1,2 and 3 suggesting additional roles during erythroid maturation. In contrast, TFs that play critical roles in HSC or in promoting non-erythroid lineages (e.g. ETV6, ERG, PU.1, FLI1, RUNX1) are expressed at higher levels at early stages (days 0-4), consistent with lineage priming, and gradually decrease as the cells progressively commit towards the erythroid lineage. Thus, our differentiation system faithfully mimics the dynamics of erythropoiesis from early to late stages.

### Major Discrepancies in Protein versus mRNA Abundances for Master Regulators of Hematopoiesis and Erythropoiesis

Having quantified protein and mRNA abundances simultaneously for multiple TFs (see for example GATA2 in Figure 1C,D), we explored the correlation between mRNA and protein levels. Consistent with previous studies in other cell types (Liu et al., 2016; Vogel and Marcotte, 2012), we found a good protein vs mRNA correlation in cells starting at day 2, with Spearman coefficients ranging from 0.40 to 0.56 (Figure 2A). In contrast, the correlation between mRNAs and proteins is extremely low (<0.25) in early progenitors at day 0 (Figure 2A). This finding, which may be explained by the low protein translation rate in HSCs (Liu et al., 2017; Signer et al., 2014), indicates that post-transcriptional regulation may be particularly important to determine protein abundance in stem/progenitor cells.

**Figure 2.**
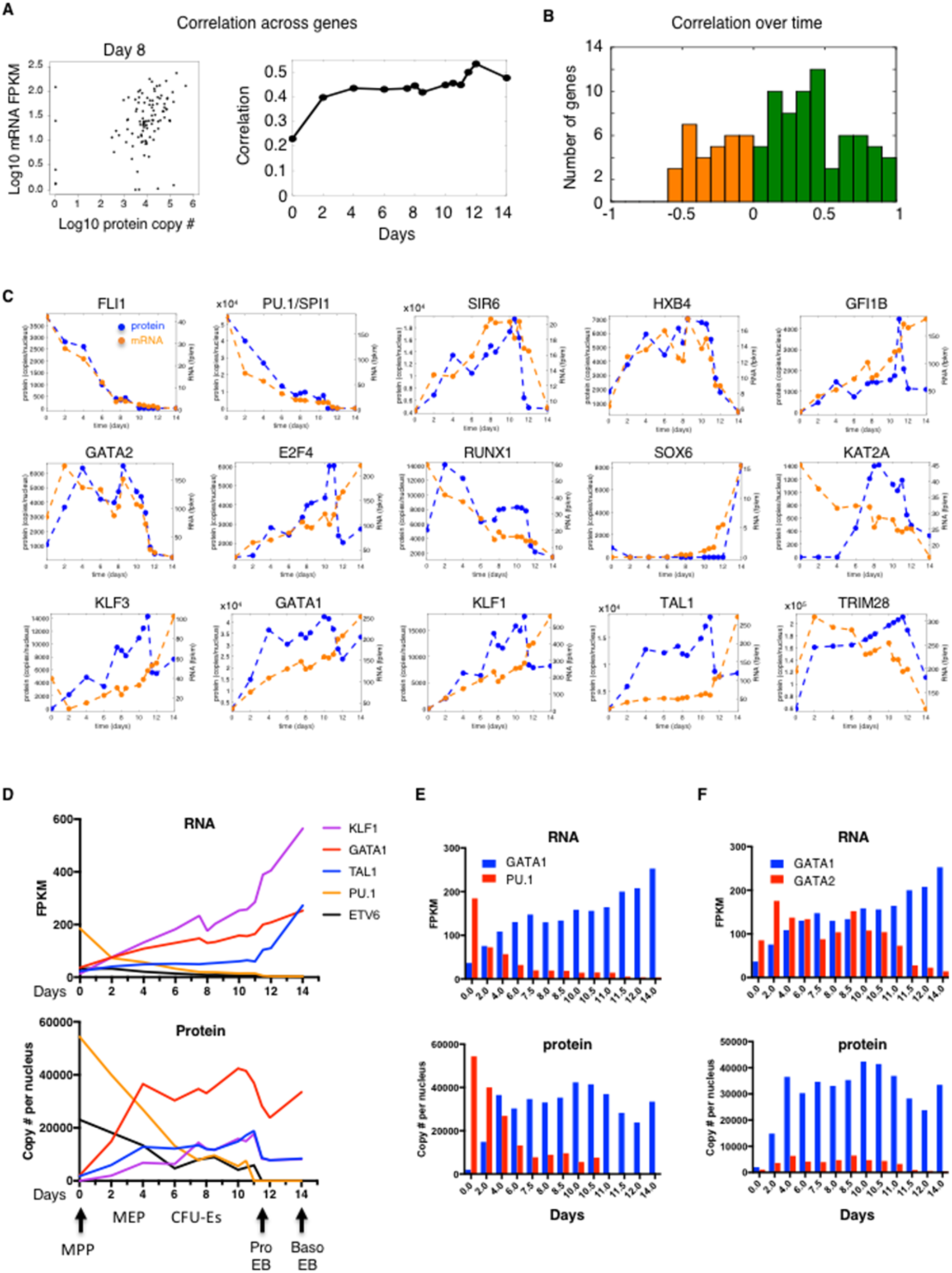
Major Discrepancies in Protein versus mRNA Abundances for Master Regulators of Hemato/Erythropoiesis. (A) Correlation between mRNA and protein abundances at the indicated days. The left panel shows a representative example of a dot plot at day 8 with each dot representing a protein-mRNA pair. The right panel shows the calculated correlation at each day. (B) Correlation between the changes in mRNA and protein levels over time during differentiation from day 0 to day 14. Positive correlations are in green. Negative correlations are in orange. (C) Protein (blue) and mRNA (orange) abundances for the indicated genes during differentiation. See Figure S5 for all 103 measured genes. (D) mRNA (top) and protein (bottom) stoichiometry for the indicated genes during differentiation. (E) mRNA (top) and protein (bottom) stoichiometry for the indicated genes during differentiation. See also Figure S4 and our Human Erythropoiesis TFs website (https://trena.systemsbiology.net/app/srm_rna_combined_v2).

Next, we measured the correlation of mRNA and protein levels over time (Figure 2B). We found that for most genes there is a positive correlation (green bars), indicating that changes in protein levels result in large part from changes in their mRNAs abundance during erythroid differentiation. However, some genes display low or negative correlations (orange bars). Among genes that display a high correlation between mRNA and protein, we found the non-erythroid TFs FLI1 and PU.1 (also called SPI1) both of which decrease gradually during erythropoiesis, but also SOX6, HOXB4 and GATA2 that increase during differentiation (Figure 2C). In contrast, other genes display major discrepancies in protein vs mRNA dynamics (e.g. KAT2A, RUNX1). Interestingly, all master regulators of erythropoiesis (i.e. GATA1, KLF1, KLF3 and TAL1) display a similar pattern of changes with protein levels increasing faster than mRNA levels until the ProEB stage (day 11.5) when they decrease by 40 to 50% even though their transcripts continue to increase (Figure 2C,D). This sharp decline in TF protein levels in the nucleus starting at the ProEB stage was also detected by iTRAQ (<u>Table S1</u>) and Western blot (Figure S1D) and coincides with a drastic change in the transcriptome (Figures 1E and S2A) as the cells enter terminal differentiation. All genes can be explored in Figure S4 or on our website (https://trena.systemsbiology.net/app/srm_rna_combined_v2).

An important aspect of transcriptional regulation is the change in relative abundances between TFs. Examining master regulators of hematopoiesis, our results reveal major differences in stoichiometry. For instance, at the transcript level, KLF1 is the most highly expressed TF in erythroid cells (Figure 2D). However, at the protein level, GATA1 is the most abundant TF with over 41,000 copies per nucleus in late CFU-E compared to fewer than 18,000 copies for KLF1, TAL1 or NFE2 (Figure 2D and Table S4). This high abundance of GATA1 protein is likely due to enhanced translational efficiency as shown by polysome profiling (Khajuria et al., 2018; Liu et al., 2017).

In addition to erythroid-specifying TFs, we also quantified TFs that are involved in stem cell maintenance (e.g. ERG, ETV6, RUNX1) and/or specification of alternate hematopoietic lineages (e.g. PU.1, FLI1, CEBPb). Interestingly, even though these non-erythroid factors gradually decrease during differentiation, they are still detectable at the protein and RNA level until late CFU-Es (Figure 2 C,D and Table S4) which suggests some degree of lineage plasticity in late progenitors. In particular, we examined the relative levels of two antagonist TFs GATA1 and PU.1, which promote mutually exclusive hematopoietic lineages (Huang et al., 2007). As expected, we observed a gradual change in the GATA1/PU.1 ratio, at both the RNA and protein levels (Figure 2E) showing that for some factors, RNA levels can be used as a surrogate for protein levels. Another TF switch necessary for erythroid maturation is the GATA1/GATA2 switch (Katsumura et al., 2017). It has been shown that during hematopoiesis GATA2 is expressed earlier than GATA1. As erythroid differentiation proceeds, GATA1 progressively represses GATA2 transcription such that when the cells reach the ProEB stage, GATA1 replaces GATA2 on target genes (Huang et al., 2016). It is currently believed that this exchange of GATA2 for GATA1 genomic binding is mediated through a reversal of the GATA2/GATA1 ratio with GATA1 becoming more abundant. Interestingly, even though this hypothesis is consistent with our RNA-based data (Figure 2F <u>top</u>), it is not supported by the proteomic data that shows GATA1 protein abundance largely exceeding that of GATA2 from the earliest CFU-E stage (Figure 2F <u>bottom</u>). Thus, it is highly unlikely that the GATA2/GATA1 switch is mediated through a reversal of GATA1 and GATA2 protein abundances as previously proposed. Rather, our proteomic data (Figure 2F) together with ChIP-seq data (Huang et al., 2016) suggest that GATA2 binds to specific genomic sites even in the presence of an excess of GATA1 and that the switch to GATA1 genomic binding at GATA2 sites occurs only upon the decrease in GATA2 protein levels at day 11.5 (ProEB stage). Notably, our finding of a high GATA1/GATA2 protein abundance ratio is consistent with previous data showing both a higher translational efficiency and a higher protein stability of GATA1 compared to GATA2 (Khajuria et al., 2018; Lurie et al., 2008; Minegishi et al., 2005). Thus, while in some cases, transcripts can be used as a surrogate for proteins, in other cases, they cannot, strongly emphasizing the importance of direct protein quantification.

### Quantitative Gene Regulatory Model of Erythroid Lineage Commitment

Gene Regulatory Networks (GRNs) provide a useful way to represent complex regulatory relationships that underlie important processes such as cell fate decisions (Dore and Crispino, 2011; Gottgens, 2015; Novershtern et al., 2011; Rothenberg, 2019). However, because of the lack of quantitative proteomic data on TFs, it has not been feasible to build networks that integrate quantitative changes in TF protein levels. Having quantified hematopoietic and erythropoietic TFs at multiple sequential time points, we sought to build a temporal GRN that integrates quantitative changes in protein and mRNA abundances of key TFs (Figure 3). We used both “core” hematopoietic factors with known functions in cell fate decisions (e.g. PU.1, GATA1, GATA2, FLI1, KLF1) and other factors (e.g. ERG, E2F4, KLF3, HOXB4), for a total of 14 genes. Focusing on transcriptional regulation, mathematical modeling with differential equations was used to explain the observed mRNA trajectory of each gene as a function of the observed protein trajectories of candidate regulators. The model is quantitative in that it estimates the relative contribution(s) of distinct TFs to the regulation of their respective targets (represented as transparency of the links - also see Figure 4A). Unique to our model, the parameters that quantify regulatory strength are linked to the absolute abundance of the proteins. This allows us to discriminate, for example, between a strong activator at low abundance and a weak activator at high abundance (represented as the thickness of the links). First, we found that the model correctly recapitulates the sequential cross-antagonisms between LS-TFs that underlie cell fate decisions, with the GATA1:PU.1 antagonism that regulates erythroid vs myeloid lineage choice (Huang et al., 2007) being detected first (day 0) followed by the KLF1:FLI1 antagonism (day 2) that regulates a subsequent erythroid vs megakaryocyte fate (Frontelo et al., 2007; Palii et al., 2019) (Figure 3B,C,D and <u>Movie S1</u>). Strikingly, these cross-antagonisms are not “equilibrated”, with for instance the repression of PU.1 by GATA1 being 3 times stronger than the reverse reaction at day 0, and 19 times stronger at day 4 (Figure 3C). This quantitative imbalance has not been described in previous network models and may reflect an early (but reversible) bias towards the erythroid lineage. Most importantly, our finding that antagonistic relationships between TFs are quantitatively imbalanced provides a mechanism to explain the observed instability of multipotent progenitors that do not exist as stable states but rather as constantly changing entities (Laurenti and Gottgens, 2018). Similarly, our model revealed the dynamics of the FLI1:KLF1 cross-antagonism that starts at day 2, decreases at day6, and finally disappears at day 8 (Figure 3D). Thus, our proteomic-based network was able for the first time to capture the temporal and quantitative aspects of gene regulation to model the dynamics of erythroid lineage commitment. To facilitate visualisation of the network, an interactive version is provided in BioTapestry format (Paquette et al., 2016) at: http://grns.biotapestry.org/HumanErythropoiesisGRN/.

**Figure 3.**
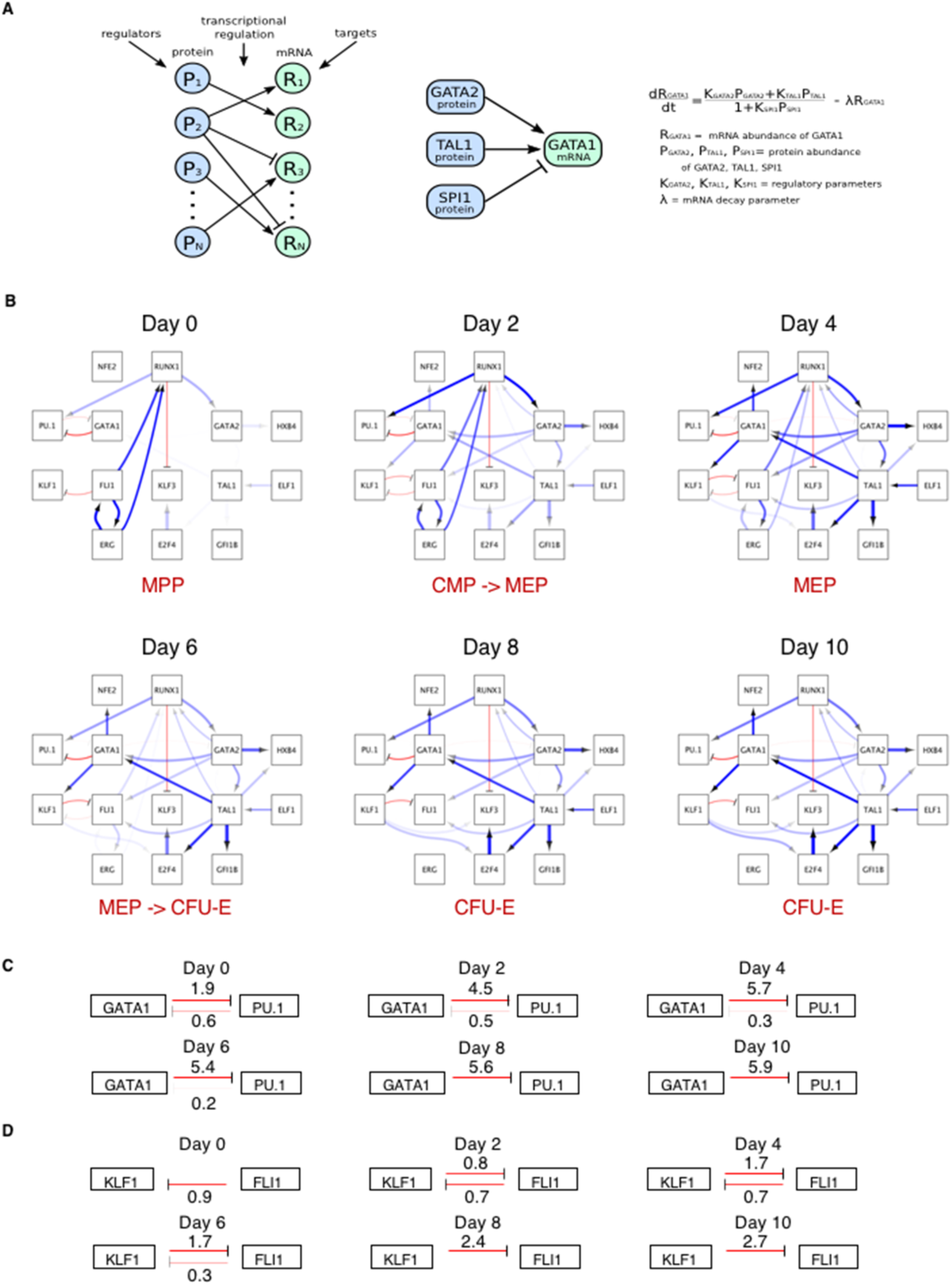
Quantitative Gene Regulatory Network of Erythroid Commitment. (A) Modeling approach: mRNA dynamics are modeled depending on regulator protein abundances. Transcriptional activation is represented as a ratio of two linear functions, corresponding to activation and repression, with constant rates multiplied by protein abundances. mRNA degradation is modelled as a linear decay. (B) Quantitative Network diagrams depicting dynamic changes in transcriptional regulation of erythroid commitment. Blue and red links indicate activation and repression, respectively. Link transparency indicates the relative contribution of a TF to the regulation of its targets (more transparent links represent weaker effects, depending on regulatory parameter and regulator abundance). Link thickness indicates the regulatory effect per TF protein molecule (thicker links indicate greater regulatory effect per TF protein molecule). The network is also available in BioTapestry format (Paquette et al., 2016) at http://grns.biotapestry.org/HumanErythropoiesisGRN/ (C,D) Quantitative imbalance in the GATA1-PU1 (C) and KLF1-FLI1 (D) cross-antagonistic relationships over time. Excerpt from the network in (B). The strengths of the regulatory relationships are indicated at each time point. Regulatory strength or transparency score was determined from the parameters Ki and Ri as log2(1+Ki*Ri). See also Movie S1.

**Figure 4.**
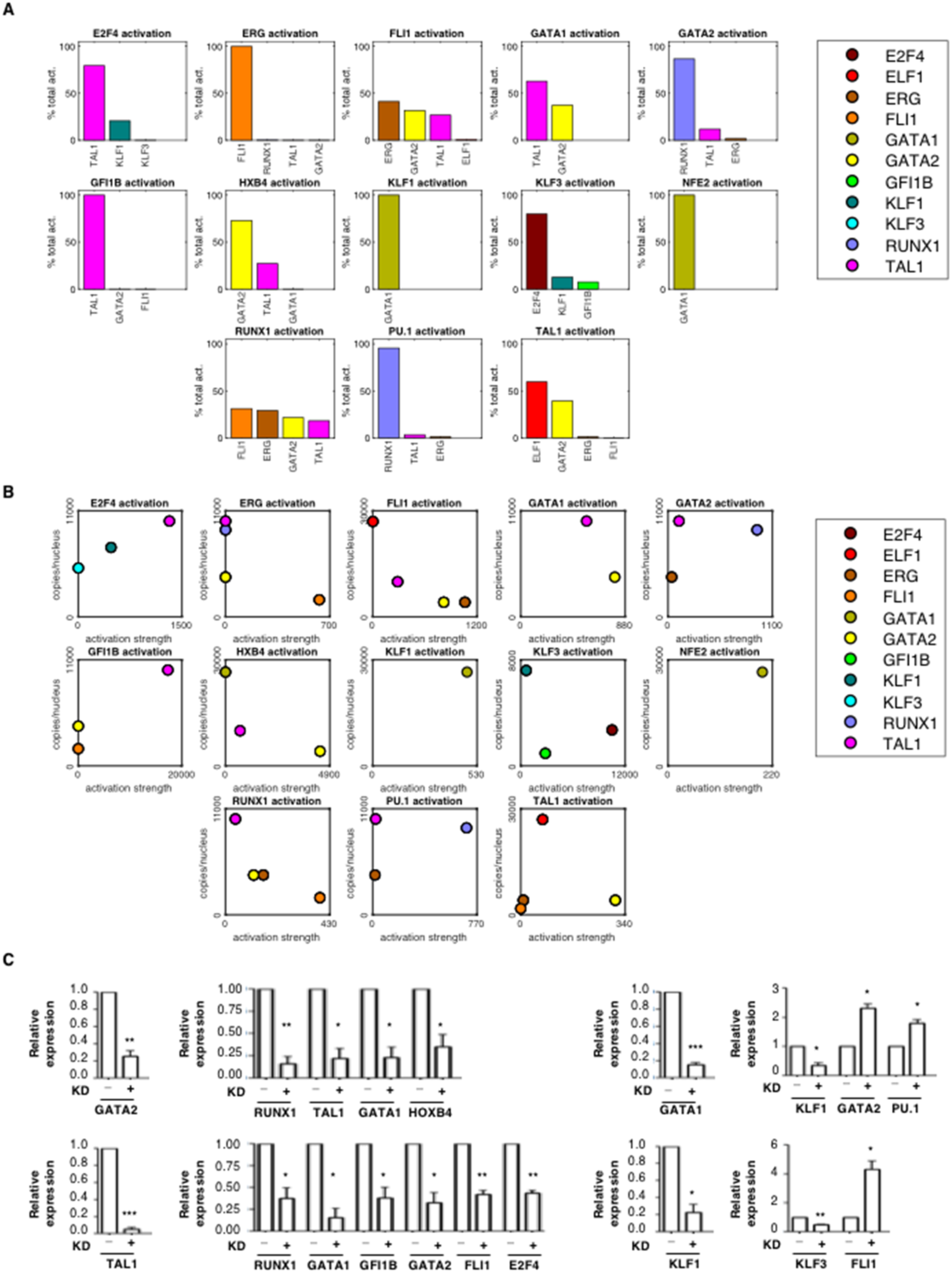
Quantitative Regulatory Relationships. (A) Relative contribution of each TF to the activation of its targets within the GRN. The average activation of a regulator X on target gene Y is computed as A_X_*K_XY_, where A_X_ is the average protein expression of gene X over the days 0, 2, 4, 6, 8 and 10, and K_XY_ is the GRN’s regulatory parameter quantifying its effect on gene Y. The activation percentage is A_X_*K_XY_ divided by the sum of average activations from all activators of Y. (B) The activation strength of each TF for its target gene is plotted relative to its abundance. For each gene Y in the GRN, the average protein abundance A_X_ and regulatory parameter K_XY_ are plotted for all activators X. (C) The knockdowns of GATA1, GATA2, TAL1 and KLF1 were induced separately by lentiviral delivery of shRNA in cells at day 8. Expression of putative target genes was tested by qRT-PCR after 48h. Expression is shown relative to GAPDH as mean +/-standard error of the mean (SEM). n=3. Two-tailed t-test: * p < 0.05. **p < 0.01. *** p < 0.001.

In addition to TF cross-antagonisms, the model confirmed a number of previously known regulatory links including the GATA1/GATA2 regulatory loop with GATA2 activating GATA1 and GATA1 progressively repressing GATA2 (Katsumura et al., 2017), the ERG-mediated activation of RUNX1 (Taoudi et al., 2011), the TAL1-mediated activation of GATA1, GFI1b and FLI1 (Gottgens, 2015), the KLF1-mediated activation of KLF3 (Ilsley et al., 2017), the GATA2-mediated activation of HOXB4 (Fujiwara et al., 2012) and others (Figure 3B). Furthermore, the model identified novel putative links such as TAL1- and KLF1-mediated activation of E2F4, E2F4-mediated activation of KLF3, TAL1-mediated activation of HOXB4 and others. These novel regulatory links are likely important for erythropoiesis. For instance, E2F4 has been shown to promote erythropoiesis (Kinross et al., 2006) and our results suggest that this may be mediated at least partially through the activation of KLF3. Most importantly, the model offers a temporal and quantitative view of these regulatory relationships. For instance, we found that over the entire time course, the contribution of TAL1 to the activation of the GATA1 gene is greater than the contribution of GATA2 (with GATA2 being more important at early stages) (Figure 4A). Interestingly, this is not because TAL1 is a stronger activator of GATA1 but instead, this is due to the fact that the TAL1 protein is on average 3 times more abundant than the GATA2 protein (Figure 4B). Thus, the model is able for the first time to quantitatively dissect the relative contributions of different factors to the regulation of their target genes.

To validate our network model, we induced the knockdown of four different TFs (i.e. GATA2, GATA1, TAL1 and KLF1), followed by qRT-PCR assessment of their target genes. All regulatory relationships tested were validated, including “activating” links (e.g. GATA2-mediated activation of RUNX1) and “repressive” links (e.g. GATA1-mediated repression of PU.1) (Figure 4C).

An important aspect of our model is that it was built without requiring data on TF genomic binding. Thus, the links within the model could reflect direct or indirect regulation. To estimate the extent to which the identified gene regulatory relationships are mediated through direct TF binding, we re-analyzed previously published ChIP-seq datasets that were available for 8 out of the 14 tested TFs in HSPCs and/or ProEBs (Methods). We found that 82% of the links for which ChIP-seq data was available can be explained by direct TF binding to either the gene promoter or an associated enhancer.

In summary, we established the first GRN that integrates quantitative changes in mRNA and protein abundance to model erythroid lineage commitment. The model correctly recapitulated known regulatory relationships and identified new links while providing information on the relative contribution of different TFs to the regulation of their targets. Most importantly, incorporation of protein concentration (i.e. stoichiometry) revealed quantitative imbalances in TFs cross-antagonistic relationships that may underlie hematopoietic progenitors’ instability and drive cell lineage commitment.

### Co-repressors are Present in Large Excess while Co-activators are Limiting Compared to TFs in the Nucleus

The function of a TF is defined by its capacity to recruit cofactors (co-activators and/or co-repressors) to target genes (Reiter et al., 2017). Thus, the function of a TF depends on the availability of cofactors in the nucleus. Yet, we do not know whether cofactors are present in excess or in limiting amounts compared to TFs. To address this, we used SRM to quantify various types of cofactors, including co-activators such as histone acetyltransferases (e.g. CBP, P300, KAT2A (also called GCN5)), histone methyltransferases (e.g. MLL1, MLL3, MLL4, SETD1B, DOT1L), histone demethylases (e.g. UTX) as well as co-repressors such as histone deacetylases (e.g. HDAC1,2,3), chromatin remodeling enzymes (e.g. CHD4 (also called Mi2b) of the NuRD complex), histone methyltransferases (e.g. SETDB1), histone demethylases (e.g. LSD1), DNA methyltransferase (e.g. DNMT1) and others (e.g. ETO2, TRIM28). First, we found that co-repressors are present in excess compared to TFs with for instance CHD4 being present at > 500,000 copies per nucleus versus 42,000 copies for the most abundant TF GATA1 at day 10 (Figure 5A,C and <u>Table S4</u>). Histone deacetylases are also highly abundant (e.g. HDAC1 > 63,000 copies per nucleus at day 10). In contrast, co-activators are surprisingly scarce with the histone acetyltransferase P300 representing less than 10% of the nuclear amount of 10 TFs combined (Figure 5B) with 7,000 copies at day 10. This surprising finding reveals a vast quantitative imbalance in HDACs versus HATs in the nucleus (Figure 5C). Examining additional co-activators and co-repressors, we found that this is a general trend with co-activators being on average 100 times less abundant than co-repressors, for all factors we have examined (Figure 5C and <u>Table S4</u>) and at all time-points during hemato/erythroid differentiation (Figure 5D and Figure S5A). Strikingly, these differences in abundance exist mostly at the protein (not at the RNA) level (Figure 5D,E and Figure S5A) suggesting a post-transcriptional regulatory mechanism. Consistent with this, we found that co-activators are more sensitive to cycloheximide treatment than co-repressors (Figure S5B) suggesting that increased protein degradation could be responsible for the scarcity of co-activators in the nucleus. Thus, our quantitative measurements revealed that the nucleus is a highly repressive environment with co-activators being exceedingly rare compared to both co-repressors and TFs.

**Figure 5.**
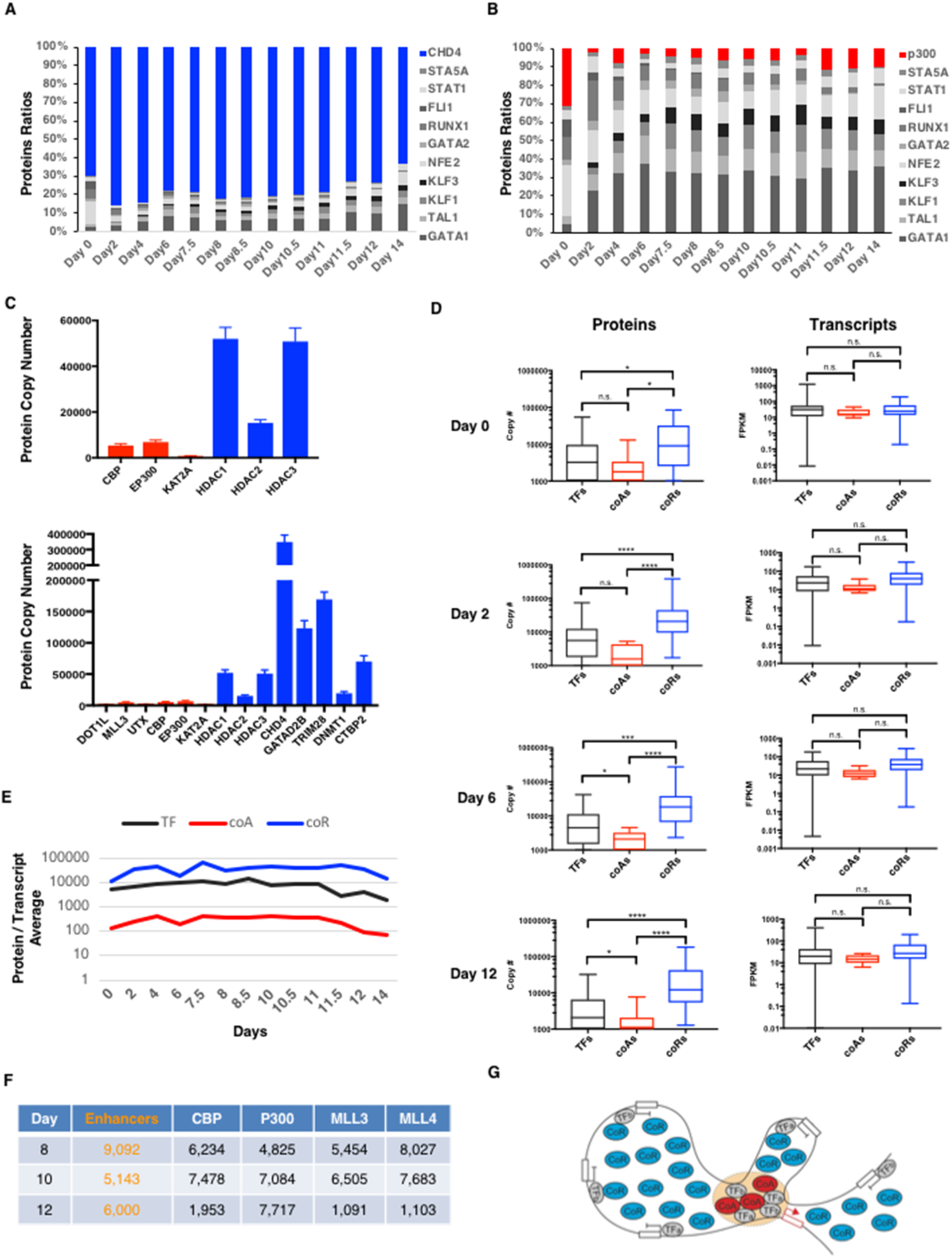
The Nucleus is a Highly Repressive Environment with a Large Excess of Co-Repressors and Limiting Amounts of Co-Activators. (A) Relative abundance of CHD4 protein compared to 10 erythroid TFs. (B) Relative abundance of P300 protein compared to 10 erythroid TFs. (C) Top panel: Quantitative imbalance between histone acetyltransferases (in red) and histone deacetylases (in blue). Bottom panel: Quantitative imbalance between co-activators (in red) and co-repressors (in blue). (D) Left panel: Box plots depicting protein abundances (in copy numbers) of transcription factors (TFs, black), co-activators (coAs, red) and co-repressors (coRs, blue) at the indicated days. Right panel: Box plots depicting mRNA abundances (in FPKM) of TFs, coAs and coRs at the indicated days. For a list of TF, coA and coR, see <u>Table S4</u>. Two-tailed t-test: n.s. (non-significant), * p < 0.05. **p < 0.01. *** p < 0.001. **** p < 0.0001. (E) Average of protein/transcript ratio for TFs, coAs and coRs at each day of differentiation. (F) Estimated number of active enhancers compared to protein copy numbers of the indicated coAs in the nucleus at the indicated days. (G) Model of gene regulation in a highly repressive nuclear environment. In this model, the same TFs are able to interact with both coAs and coRs (Lambert et al., 2018). However, due to the scarcity of coAs in the nucleus, TFs must compete to recruit them, and they do so by creating an environment that allows multiple low affinity interactions as observed on enhancers (Farley et al., 2015; Hahn, 2018). In contrast, the high abundance of coRs increases their availability, which facilitates their recruitment to genes even in case of a small number of low affinity interactions with TFs. See also Figure S5.

TFs do not work in isolation, but instead upon binding to DNA, they form highly organized structures at enhancers to facilitate the recruitment of co-activators (Catarino and Stark, 2018). Thus, we asked whether co-activators are limiting or in excess compared to active enhancers. To estimate the number of active enhancers in erythroid cells, we performed ATAC-seq (Buenrostro et al., 2013) followed by HINT-ATAC (Li et al., 2019) at three time-points to identify TFs footprints within regions of open chromatin. These regions were then intersected with predicted enhancers from the GeneHancer database (Fishilevich et al., 2017). Since enhancers are characterized by specific histone modifications (H3K27ac and H3K4me1), we compared the number of identified enhancers with the number of molecules of co-activator enzymes responsible for these histone marks, including UTX (that demethylates H3K27), CBP and P300 (that acetylate H3K27) as well as MLL3 and MLL4 (that monomethylate H3K4). Interestingly, we found that the copy number of histone-modifying enzymes is on the same order of magnitude as the number of active enhancers (Figure 5F), suggesting that enhancer formation is dependent on co-activator molecules availability in the nucleus and that enhancers must compete with each other to recruit rare and unstable co-activators in a highly repressive nuclear environment (Figure 5G).

## Discussion

In this study, by measuring absolute abundances of TFs during the course of erythropoiesis, we uncovered major discrepancies between mRNA and protein levels for master regulators of hematopoiesis. Integration of protein stoichiometry data with mRNA measurements over time allowed us to establish a dynamic GRN, which revealed quantitative imbalances in TFs cross-antagonistic relationships that underlie lineage determination. Furthermore, comparing the abundances of coactivators and corepressors in the nucleus, we made the unexpected discovery that corepressors are dramatically more abundant, which has profound implications for understanding transcriptional regulation.

A central question in biology is how TFs control gene expression programs to direct cell processes such as lineage commitment. A pre-requisite for understanding the general principles of transcription is to define the key components (i.e. transcription factors and cofactors) quantitatively in absolute terms. Yet, despite years of studies we still do not know the copy number of TFs and cofactors in the nucleus during dynamic cellular processes. We also do not know the relative abundances of coactivators, corepressors and TFs. Here, through the use of targeted MS combined with spiked-in protein-specific standards, we offer an unprecedented view of the transcriptional machinery in the nucleus, providing an abundance scale for 103 major players in transcriptional regulation, including TFs, chromatin-modifying enzymes, coactivators, corepressors and subunits of the general transcriptional machinery at different stages of differentiation from HSPCs to erythroid cells. In addition to providing a quantitative scale for TFs in the nucleus, the protein data has changed our view on key aspects of erythroid differentiation. For instance, based on mRNA measurements, it was believed that the switch that occurs at the ProEB stage between GATA2 and GATA1 genomic binding to activate an erythroid gene program was due to GATA1 becoming more abundant than GATA2 at this stage (Katsumura et al., 2017). However, our protein data show that GATA1 is more abundant than GATA2 from the early stages of hematopoiesis. Thus, in contrast to the prevailing view, GATA2 is able to bind to its target genes despite an excess of GATA1, and the GATA2/GATA1 switch in genomic binding is not mediated through an inversion in the ratio between these proteins.

Another major finding is that in the nucleus, corepressors are highly abundant (e.g. >500,000 molecules per nucleus for CHD4) whereas coactivators such as P300 or CBP are comparatively very rare (<8,000 molecules per nucleus) with TFs being present at intermediate levels. Our finding that coactivators are limiting compared to TFs is consistent with the concept of “cofactor squelching” (also called “transcription interference”)(Meyer et al., 1989) which proposes that TFs compete for a limited number of cofactors in the nucleus. The squelching model was suggested 30 years ago based on reporter assays(Meyer et al., 1989) but had remained controversial due to the lack of data on TFs vs cofactors stoichiometry in cells (Schmidt et al., 2016). Our finding that coactivators are limiting compared to TFs in the nucleus provides strong support for a model whereby TFs compete with each other to recruit a limiting number of co-activators. Furthermore, this is compatible with emerging models of enhancer function that involve multiple weak interactions between coactivators, TFs and their genomic binding sites to achieve specificity in gene regulation (Farley et al., 2015; Hahn, 2018) (Figure 5G). In this regard, it is also interesting that the number of coactivator molecules and active enhancers are roughly equivalent in the nucleus, suggesting that the formation of active enhancers may depend on coactivator availability. Although the squelching model proposes passive repression as a mechanism for genes to remain inactive due to a lack of available coactivators, the high amounts of corepressors we detected indicate that the nucleus is a highly repressive environment with the recruitment of corepressors to target genes likely facilitated by their high abundance (Figure 5G). Consistent with this, NuRD has been shown to repress transcription of fetal ß-like globin genes in adult erythroid cells (Yu et al., 2019), and to suppress transcriptional noise during lineage commitment (Burgold et al., 2019). Based on these data, we propose that restricting the abundance of coactivators in a highly repressive nuclear environment is an important yet under-appreciated mechanism for concerted gene regulation during cellular processes such as cell fate decision, by ensuring only a limited number of genes can be expressed, and thus preventing high level co-expression of lineage-specific genes in multipotent progenitors.

Cell fate decisions are thought to be mediated through competition between LS-TFs that are co-expressed in multipotent progenitors (Huang et al., 2007; Palii et al., 2019). These regulatory relationships are typically represented by GRNs wherein LS-TFs inhibit each other within a stable progenitor state (Dore and Crispino, 2011; Gottgens, 2015). However, recent single cell analyses at the RNA (Zheng et al., 2018) and protein (Palii et al., 2019) levels have suggested that hematopoietic progenitors do not exist as a stable state but instead are gradually differentiating along lineage trajectories. Thus, there are uncertainties about how well previous GRN models capture lineage commitment. A major limitation of previous network models is that they did not incorporate quantitative changes in TF protein levels and instead used mRNA as a proxy for proteins. This can negatively affect the utility of GRNs as post-transcriptional regulatory mechanisms can result in major discrepancies between RNA and protein abundances (Liu et al., 2016). In contrast, we have integrated complementary measures on mRNA and protein abundances to build a temporal GRN of erythroid lineage commitment. Our model is unique in that it is both dynamic, allowing us to capture changes in regulatory relationships over time, and quantitative as it measures the strength of each regulatory relationship and the relative contribution(s) of different TFs to the regulation of their target genes (Figures 3 and 4 and <u>Movie S1</u>). Notably our model was able to accurately recapitulate the known cross-antagonisms between LS-TFs in their correct sequential order. Most importantly, the model revealed that these cross-antagonisms are quantitatively imbalanced and that these imbalances become more pronounced with time. This suggests that lineage commitment can be quantified by measuring the imbalance between LS-TFs’ cross-antagonistic relationships. In summary, our GRN offers the first protein-based quantitative view of dynamic changes in gene regulatory relationships that underlie erythroid lineage commitment from HSPCs. We expect it will serve as a framework for integration of additional parameters such as other TFs/cofactors, post-translational modifications and/or genomic DNA binding data to allow for a more comprehensive understanding of transcriptional regulation during erythropoiesis.

Through the simultaneous measurement of RNA and proteins at multiple time points during erythropoiesis, we reveal major principles of transcriptional regulation that underlie lineage commitment. It is likely that the regulatory principles established here for erythropoiesis will be generally applicable to other cell types.

## Acknowledgements

We thank L. Tora of IGBMC (Strasbourg), D. Trono and P. Aebischer of EPFL (Lausanne) for reagents, T. Olender of OHRI for data curation, F.J. Dilworth and W.L. Stanford of OHRI for critical review of the manuscript. Library preparation and RNA sequencing were performed at the Genomics Core Facility of the Institut de Recherche Clinique de Montreal (Montreal, QC, Canada).

## Author contributions

T.J.P., J.A.R. and M.B. conceived and designed the study. C.G.P. performed cell differentiation, RNA/protein extraction and model validation under M.B. supervision. M.G. performed iTRAQ and SRM assays design and analyses under J.A.R. supervision. D.S.T. and T.J.P. performed mathematical modeling and bioinformatic analyses with some help from K.S. D.J.D. performed ATACseq experiments under J.R.H. supervision. P.S. analyzed ATACseq data. P.S. designed and implemented the TF website under N.P. supervision. W.J.R.L. designed and implemented the BioTapestry website. M.B. wrote the initial manuscript draft. M.G., D.S.T., C.G.P., J.A.R. and T.J.P. revised the manuscript. All authors discussed the results and commented on the manuscript.;

## Supplemental Figures

**Figure S1.**
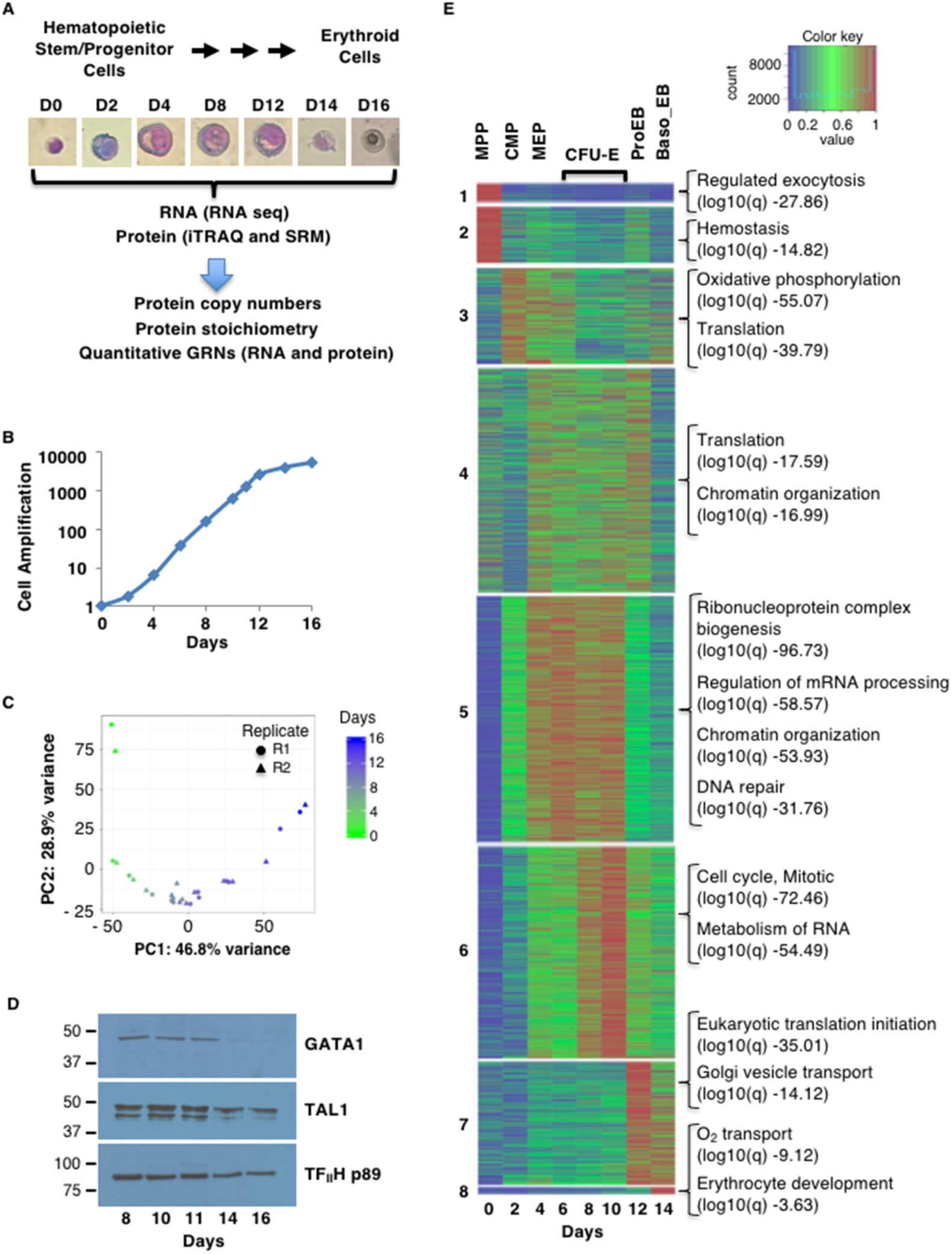
(Related to Figure 1) RNA and Protein Quantification of Transcription Factors and Co-factors during Ex Vivo Human Erythropoiesis. (A) Schematic of sample collection at different time-points followed by analyses at the RNA and protein levels. Erythroid differentiation was induced ex vivo from cord blood-derived CD34^+^ HSPCs. Giemsa-stained cells are shown at representative days (magnification 40x). (B) Cell amplification during ex vivo erythropoiesis. (C) Principal component analysis (PCA) monitoring gene expression changes over time as measured by RNAseq. (D) Western blot analysis of GATA1, TAL1 and TF_II_Hp89 proteins at the indicated days during ex vivo erythropoiesis. Molecular masses (in kDa) are indicated on the left. (E) k-means clustering analysis of iTRAQ data at different time-points. The top enriched Gene Ontology terms for each cluster are indicated.

**Figure S2.**
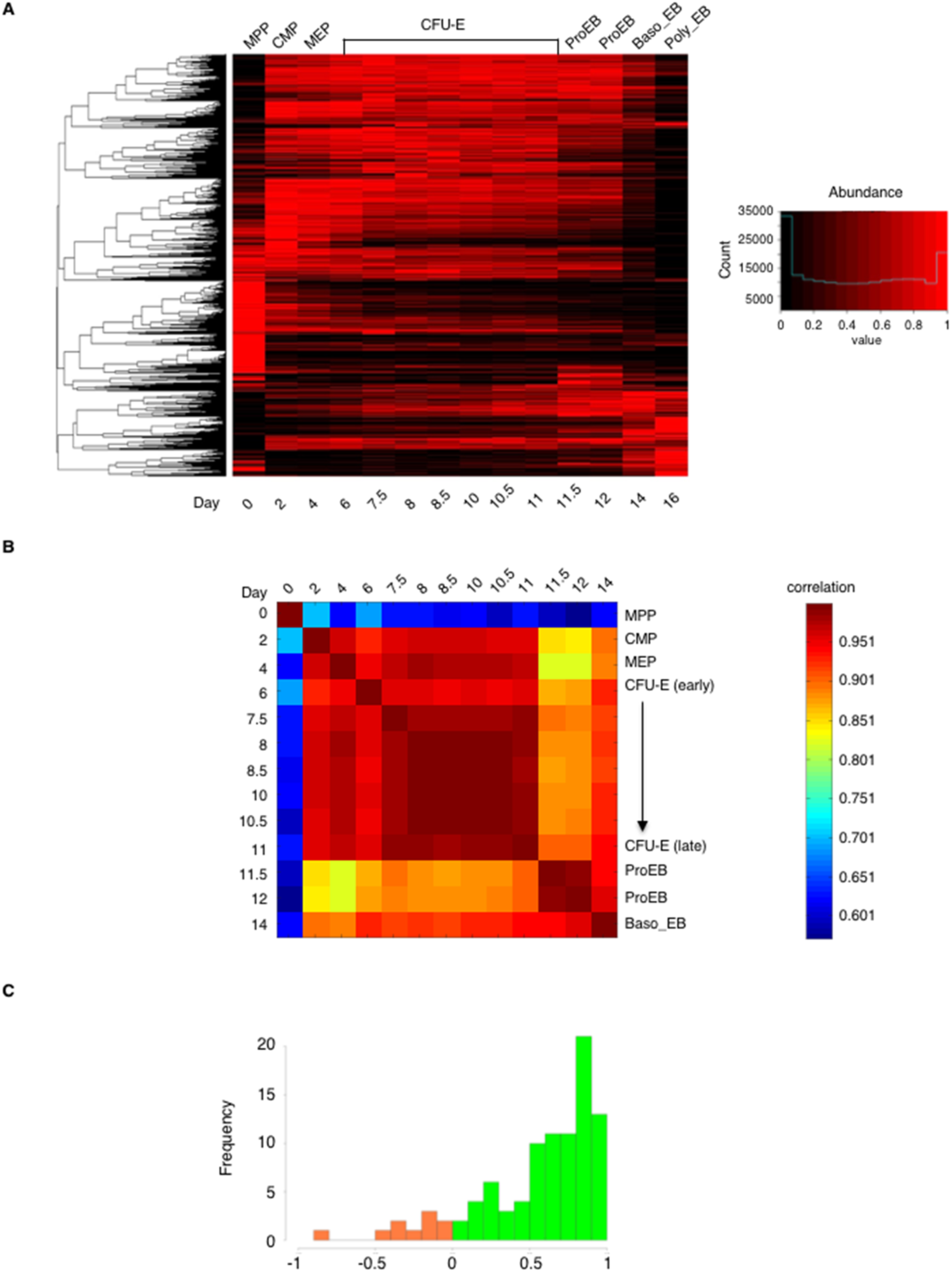
(Related to Figure 1) RNA and Protein Quantification of Transcription Factors and Co-factors during Ex Vivo Human Erythropoiesis. (A) k-means clustering analysis of normalized mRNA expression (measured by RNA-seq) at the indicated days. (B) Correlation matrix of normalized protein expression (measured by SRM) at the indicated days. (C) Correlation of protein changes over time as measured by iTRAQ and SRM. Positive correlations are in green. Negative correlations are in orange.

**Figure S3.**
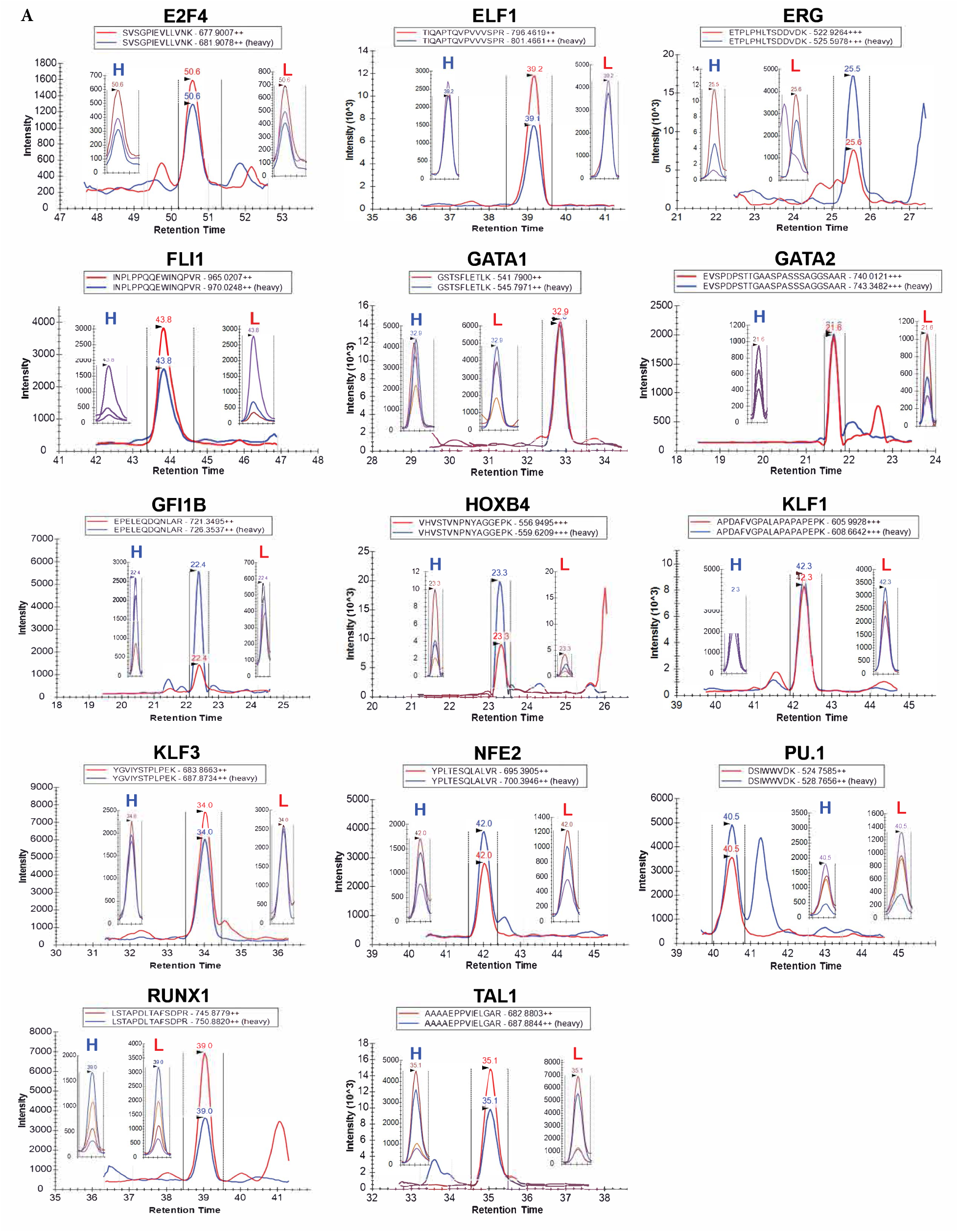
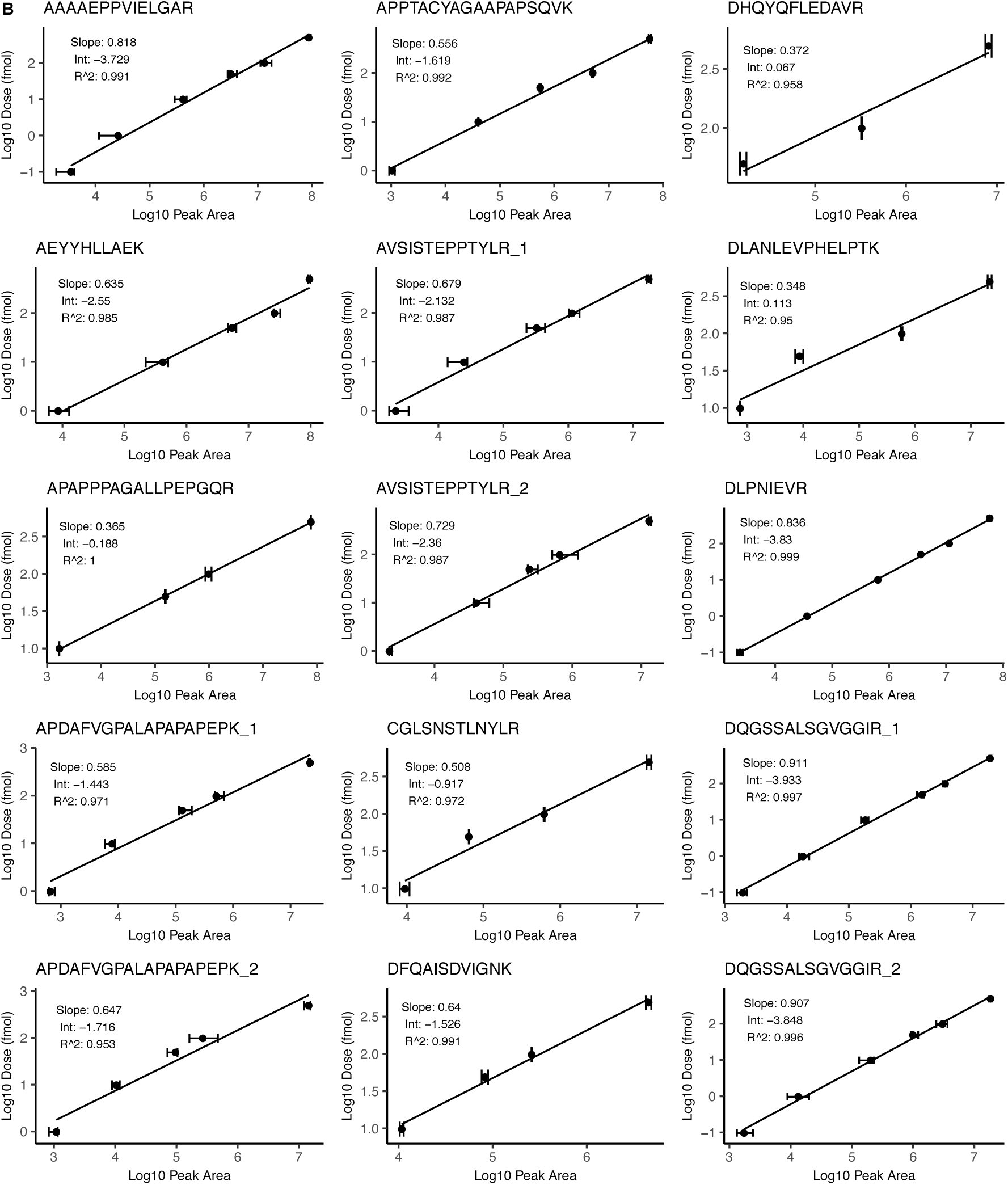
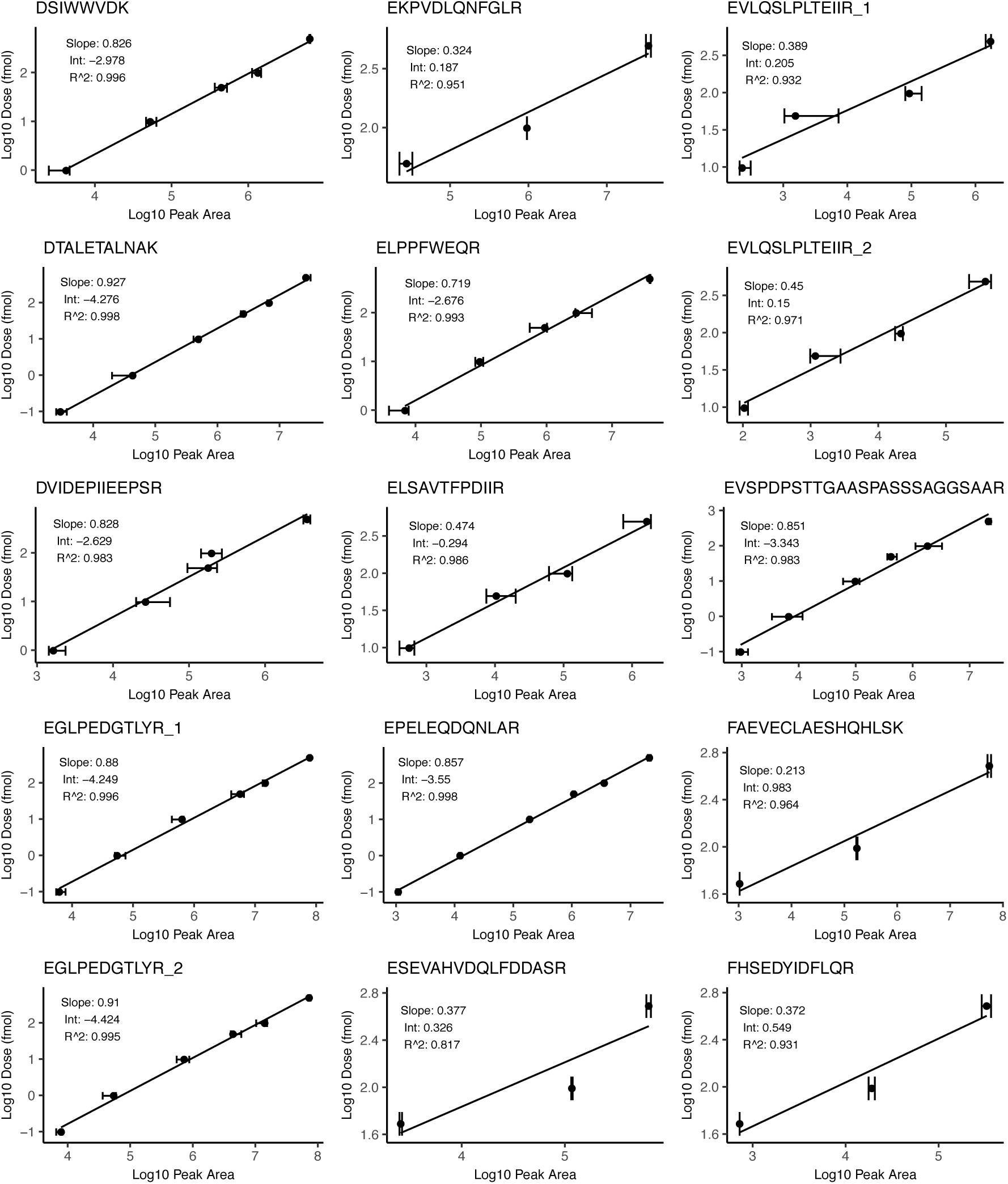
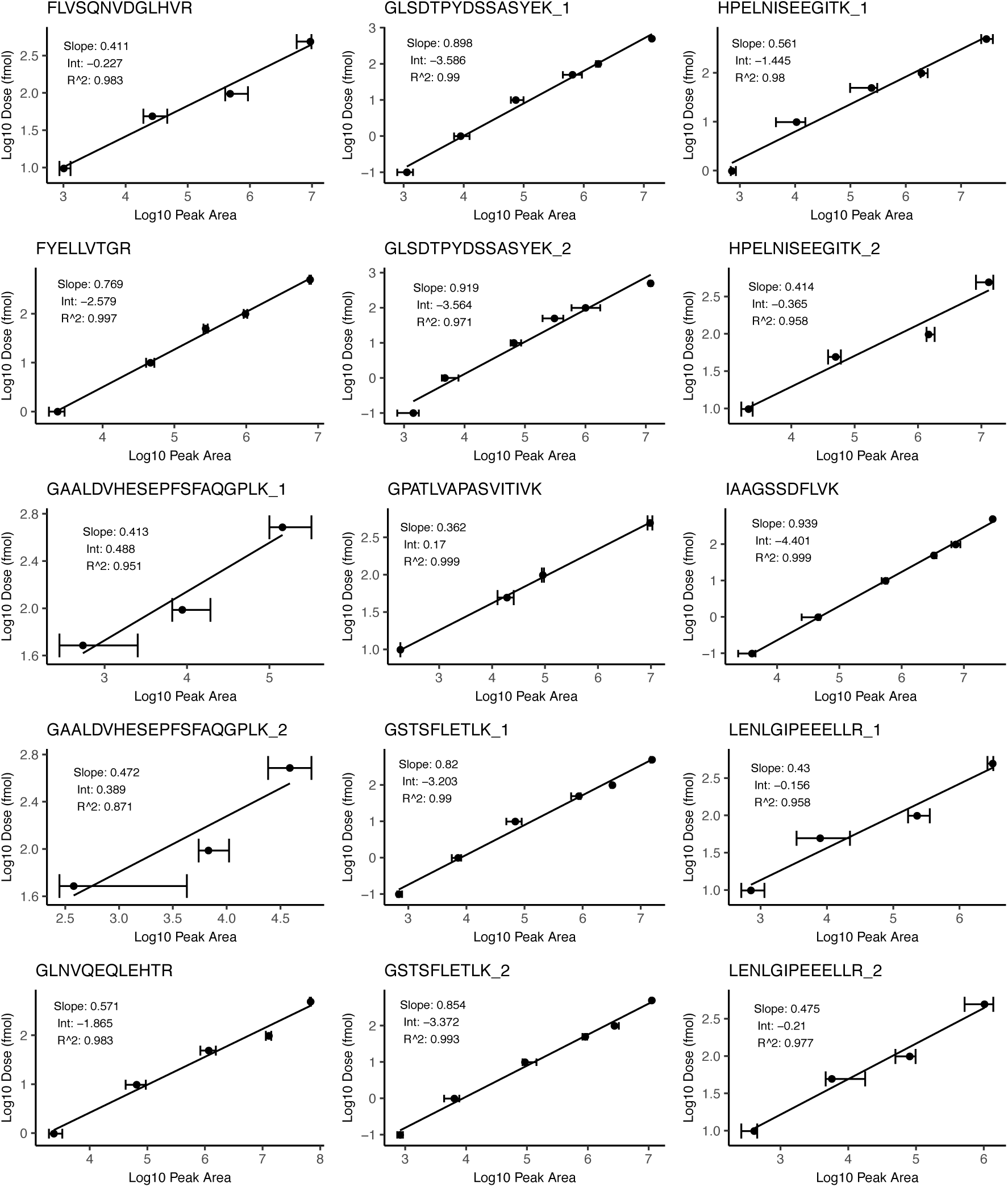
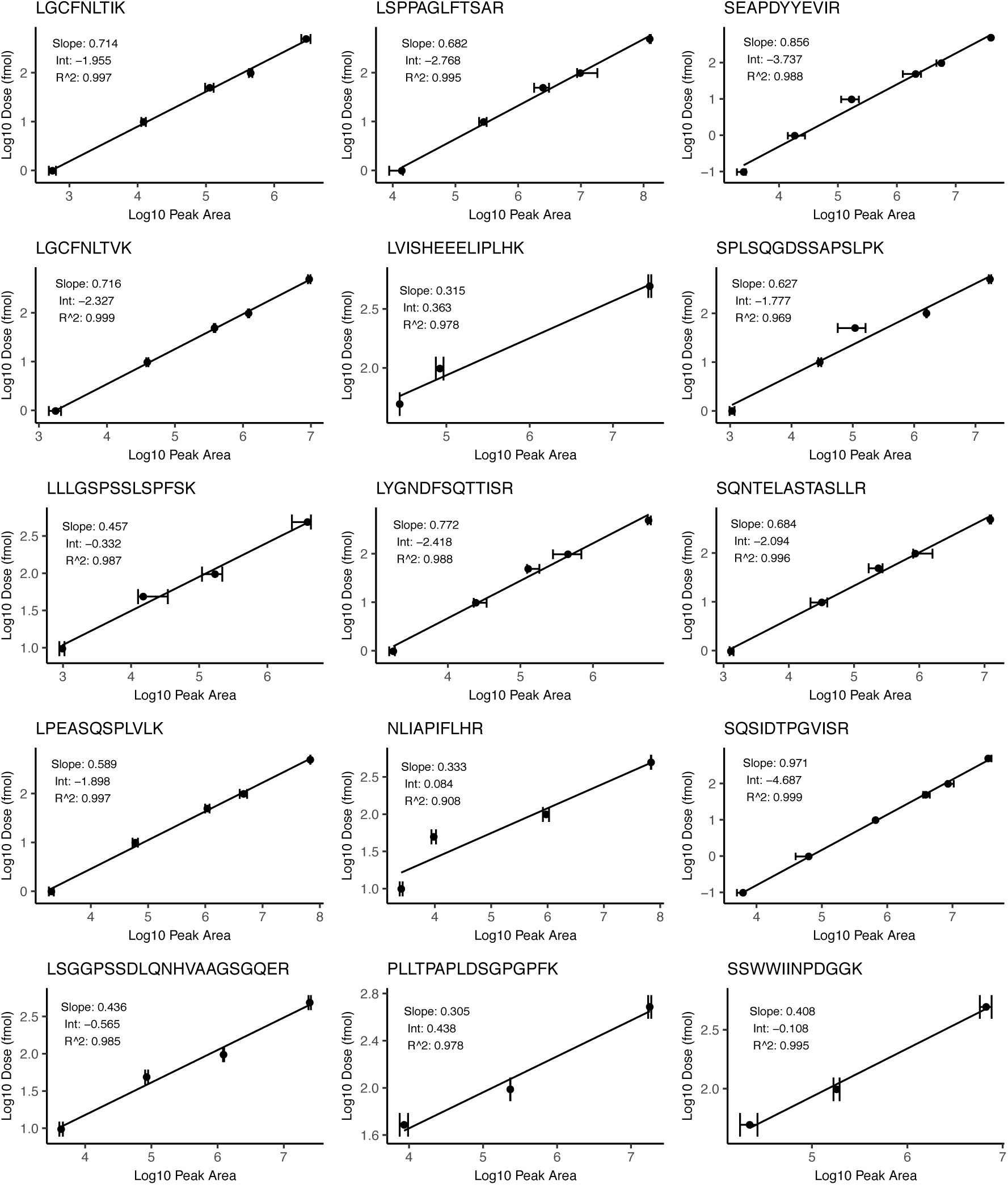
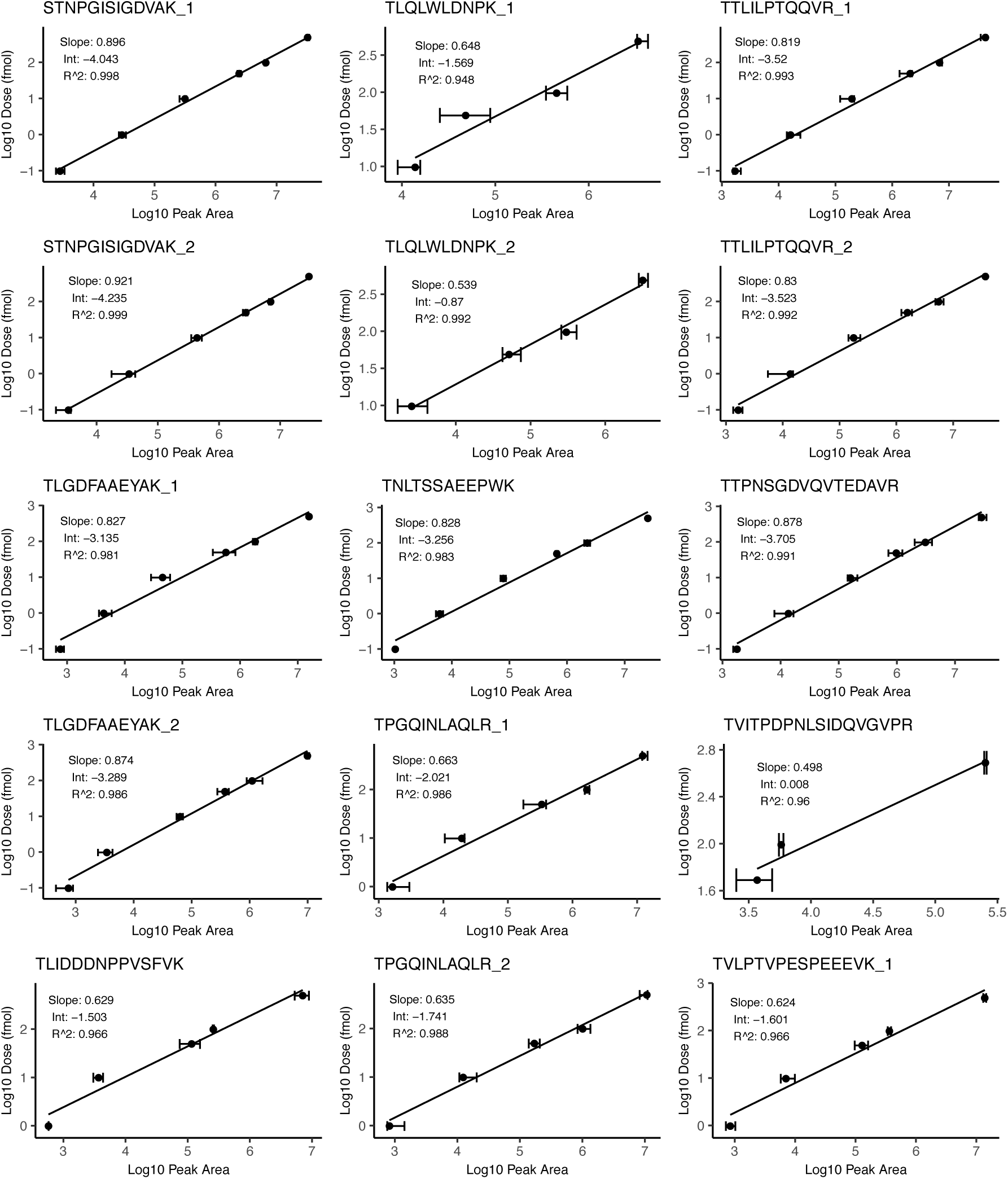
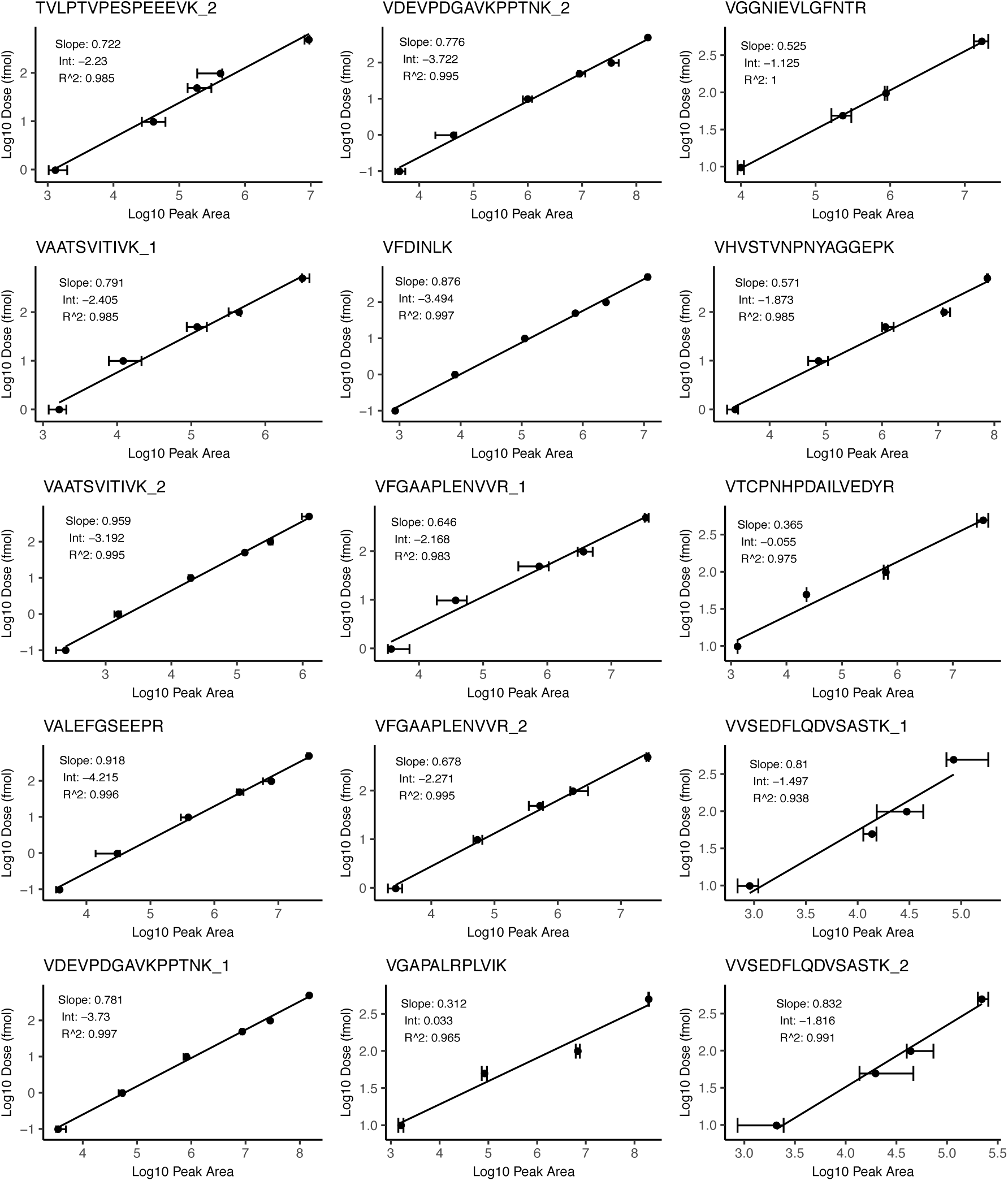
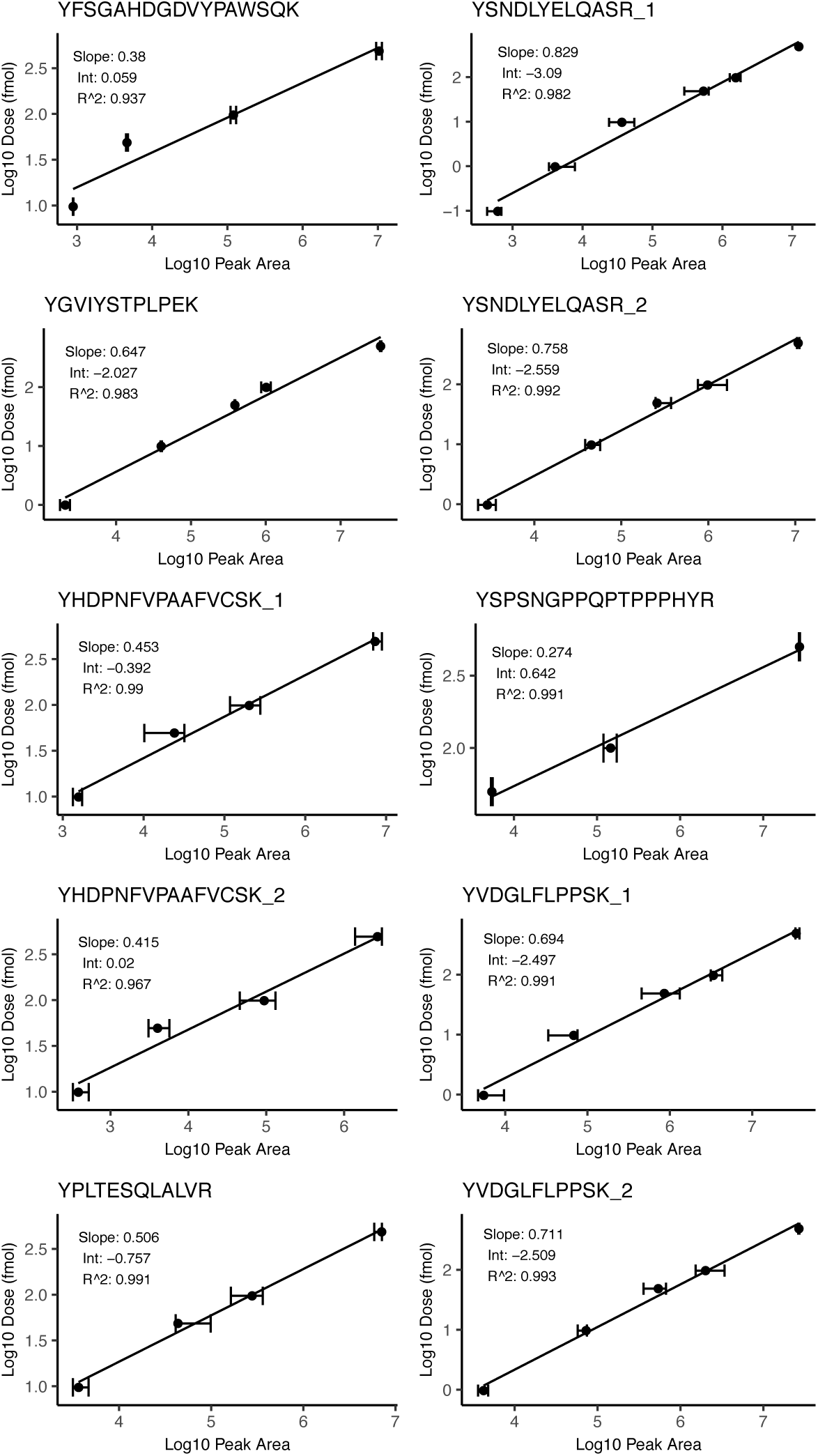
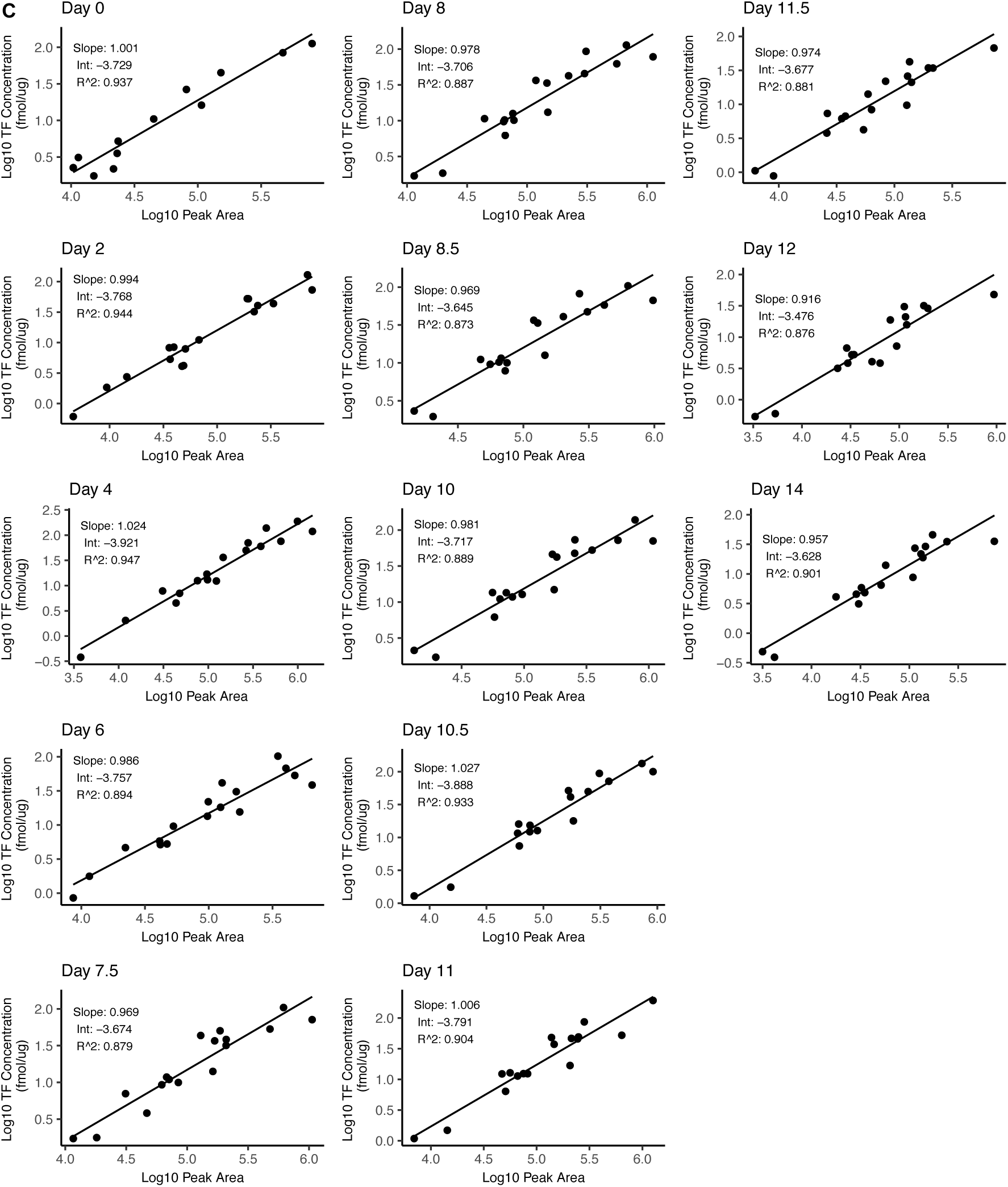
(Related to Figure 1) SRM Related Plots. (A) Representative ion chromatograms of core network transcription factors measured by SRM. The main picture shows endogenous, isotopically light-labeled peptides (red) co-eluting with exogenous SIL peptides (blue). Inset pictures show specific transitions measured for each peptide isotope (H = heavy, L = light). Images were generated in Skyline and use Savitzky-Golay smoothing. (B) Dose-response linear regression plots for all IS peptide standards (in solvent) (C) Standard curves used for label free absolute quantification. Separate curves and regression parameters were generated for each time point of differentiation.

**Figure S4.**
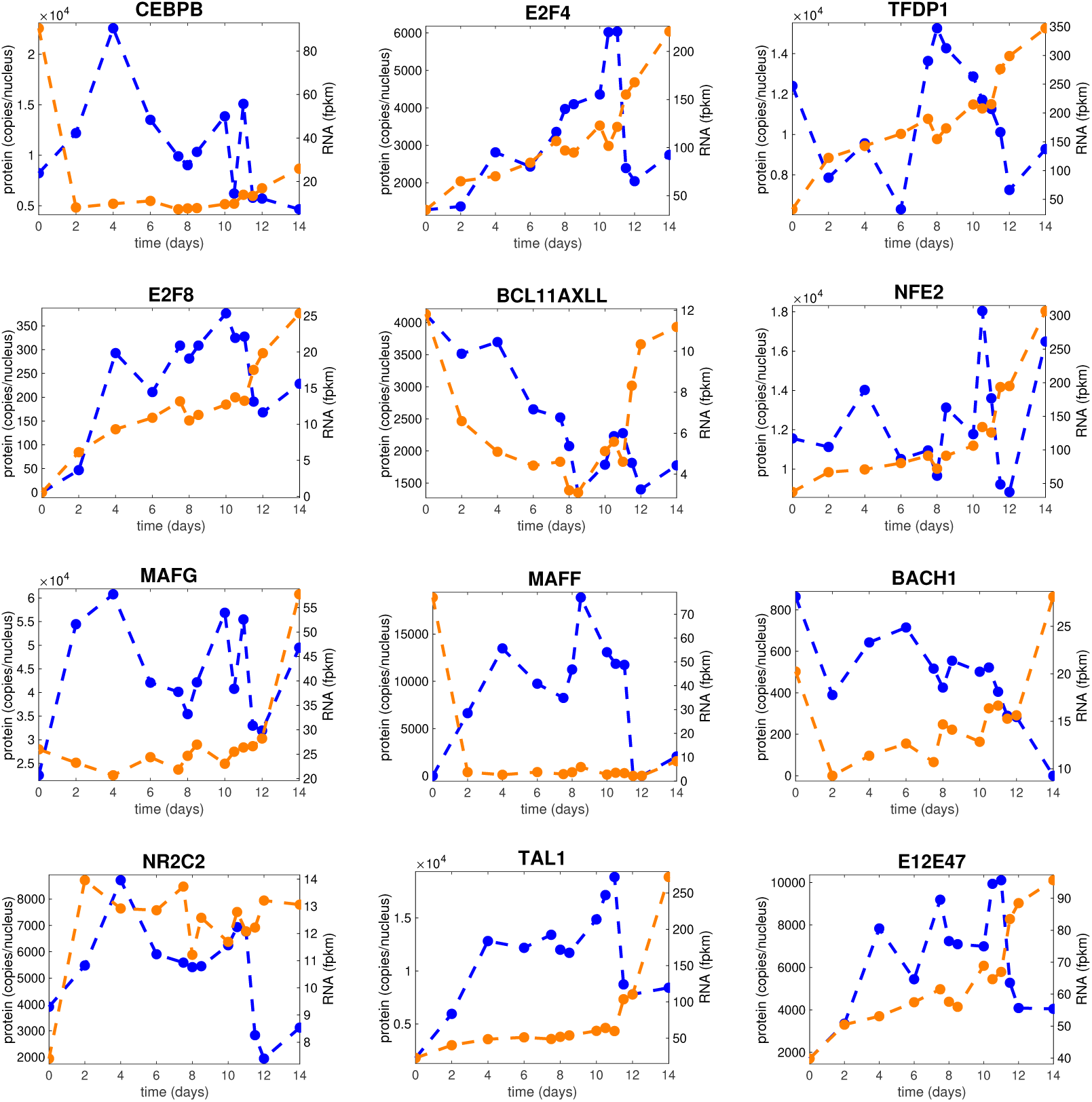
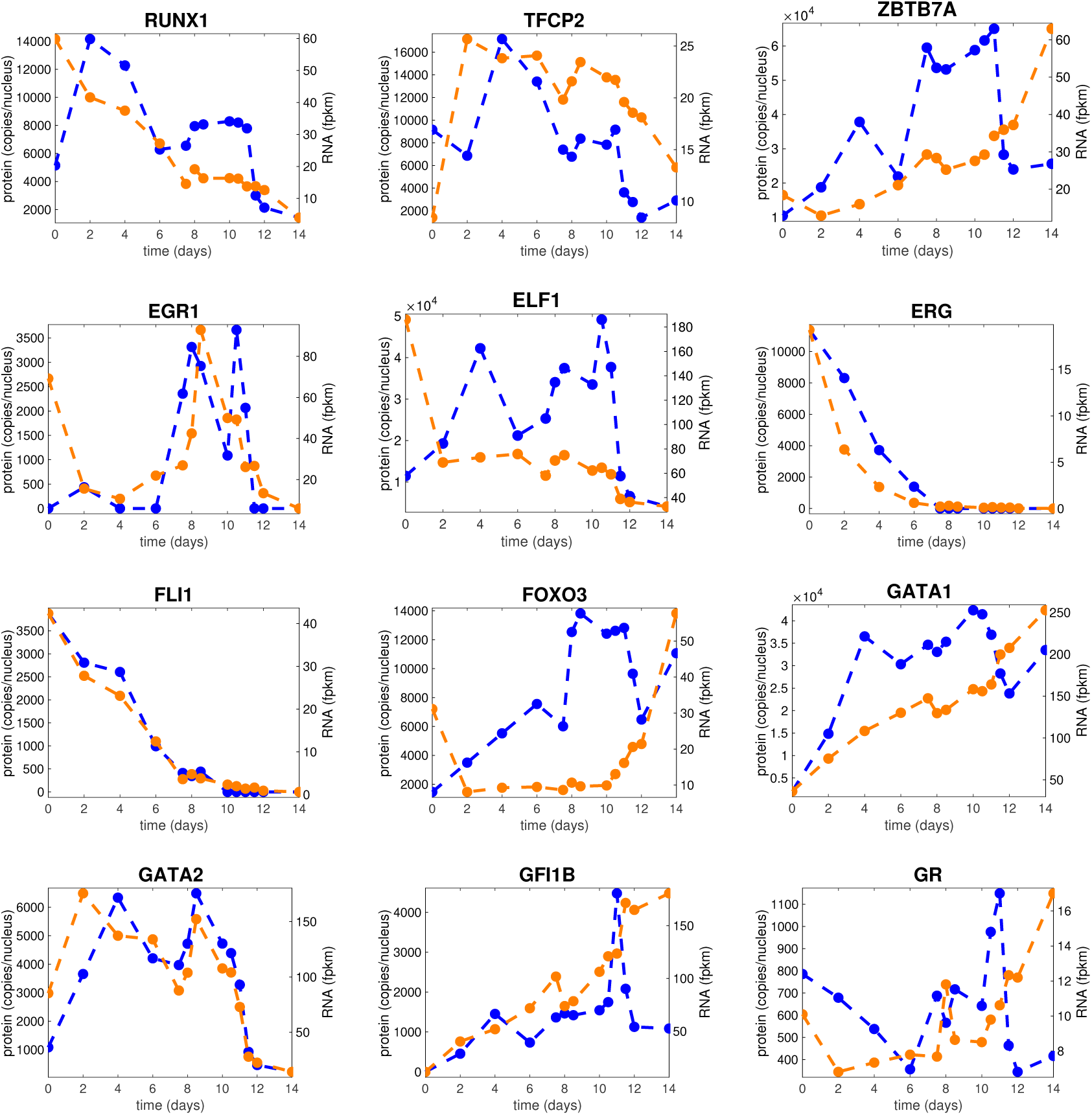
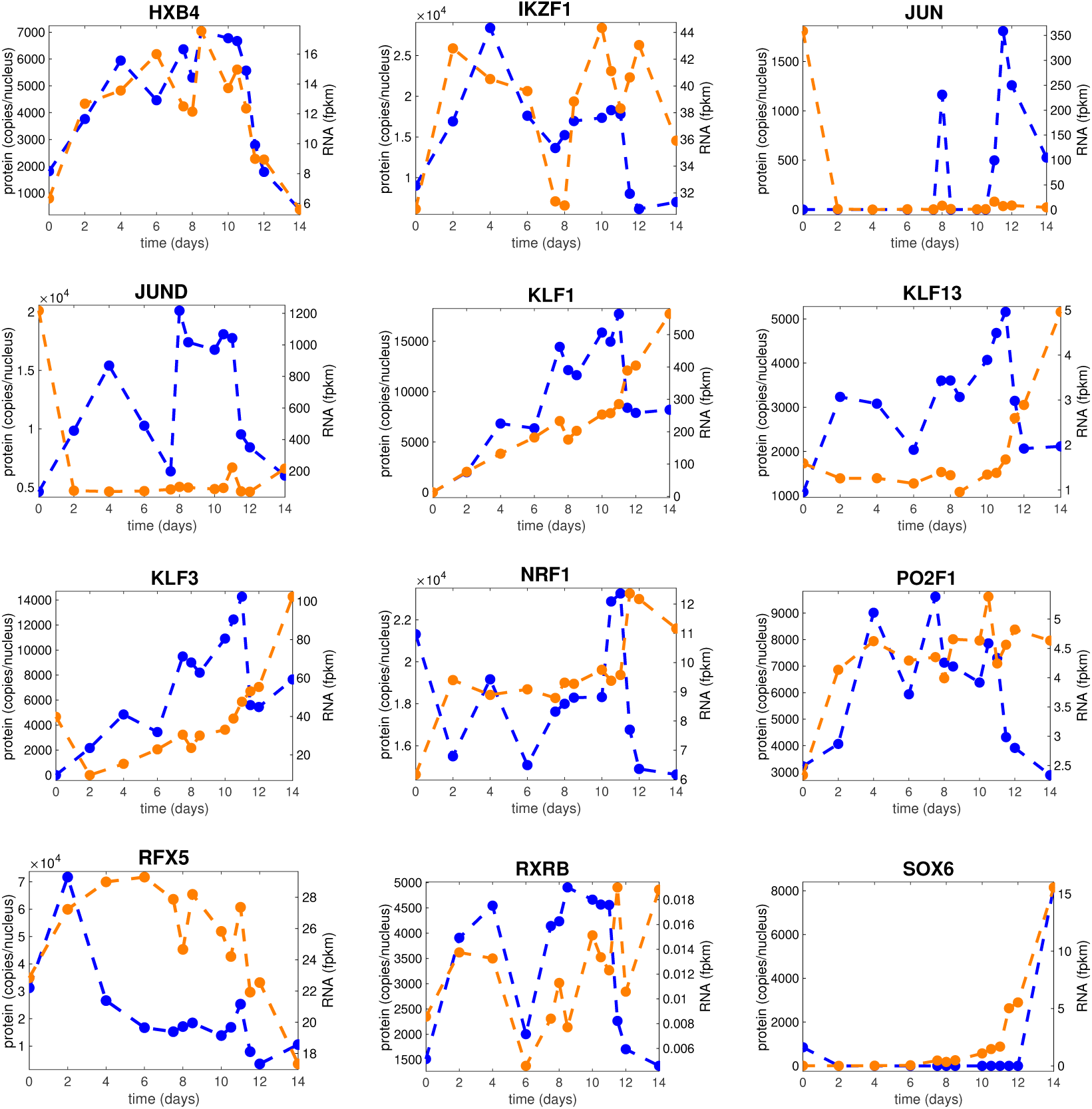
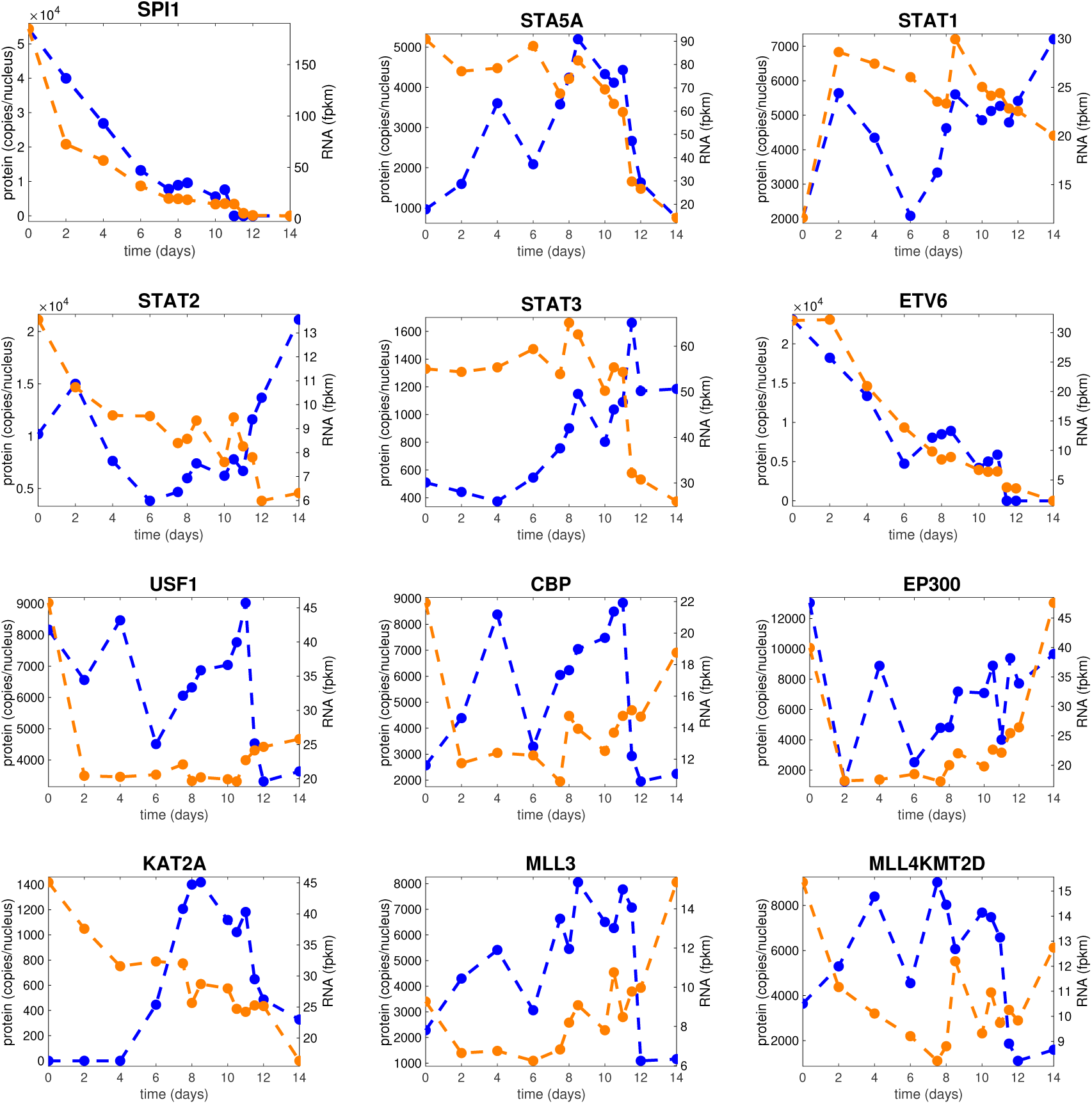
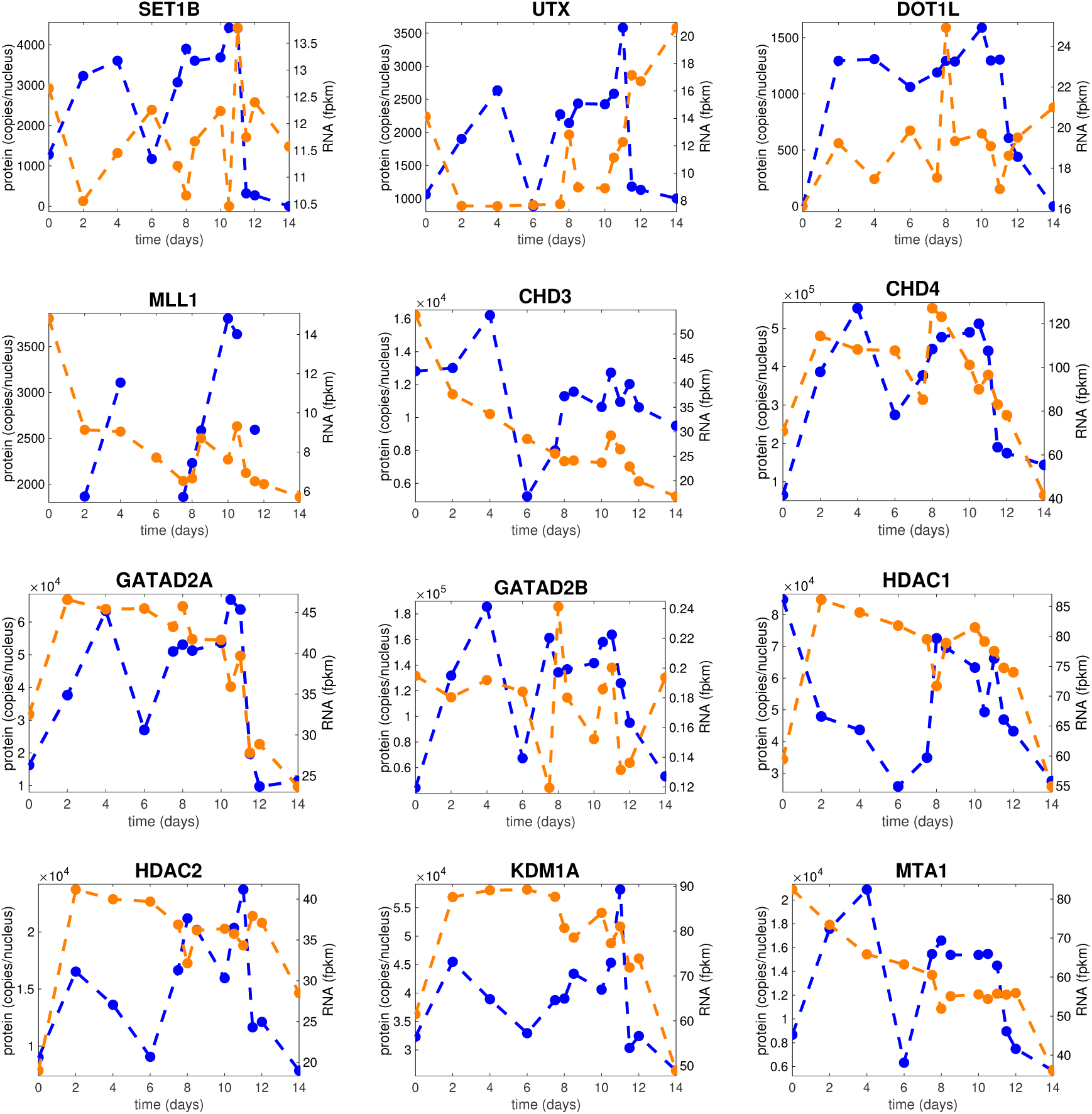
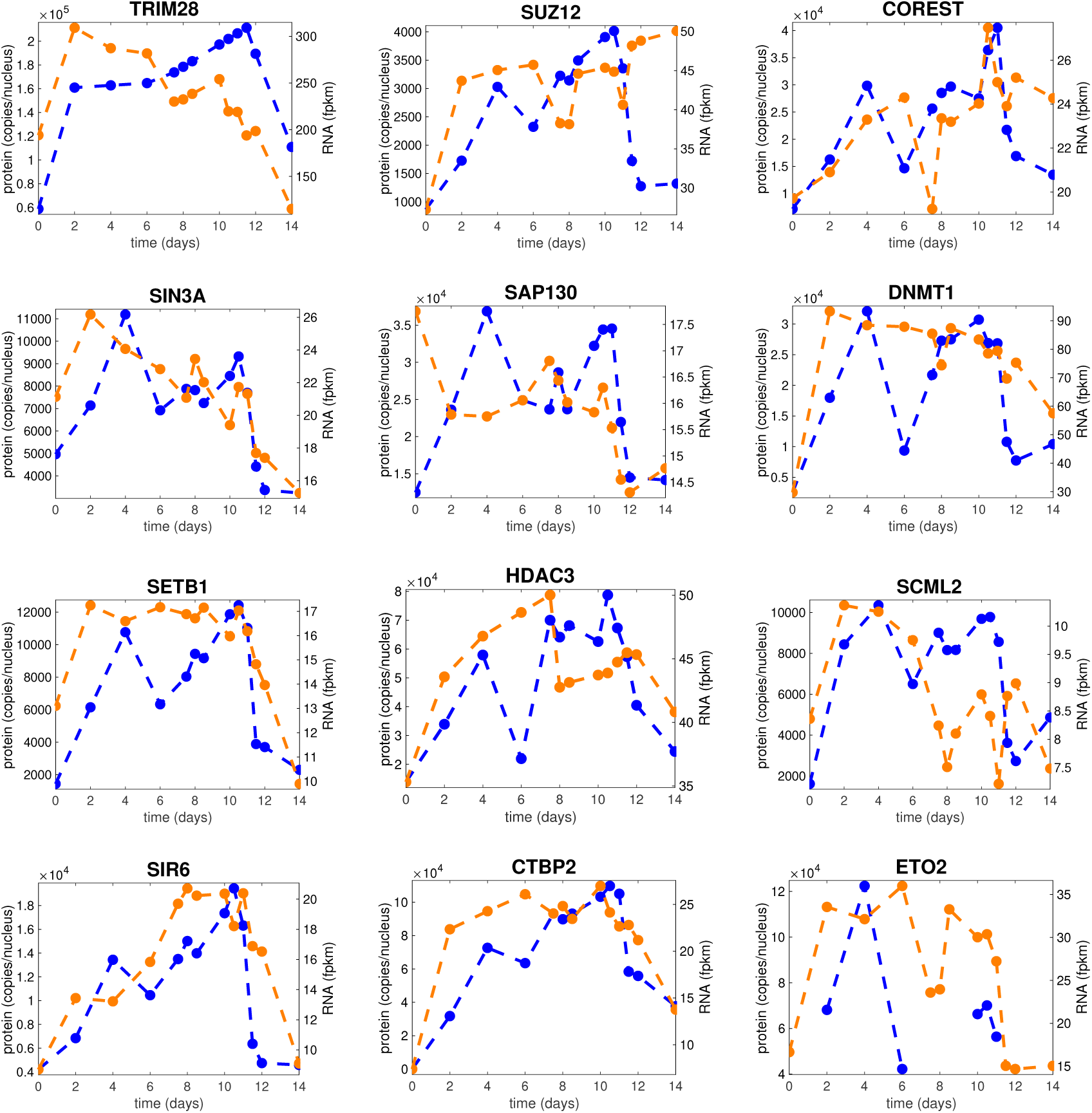
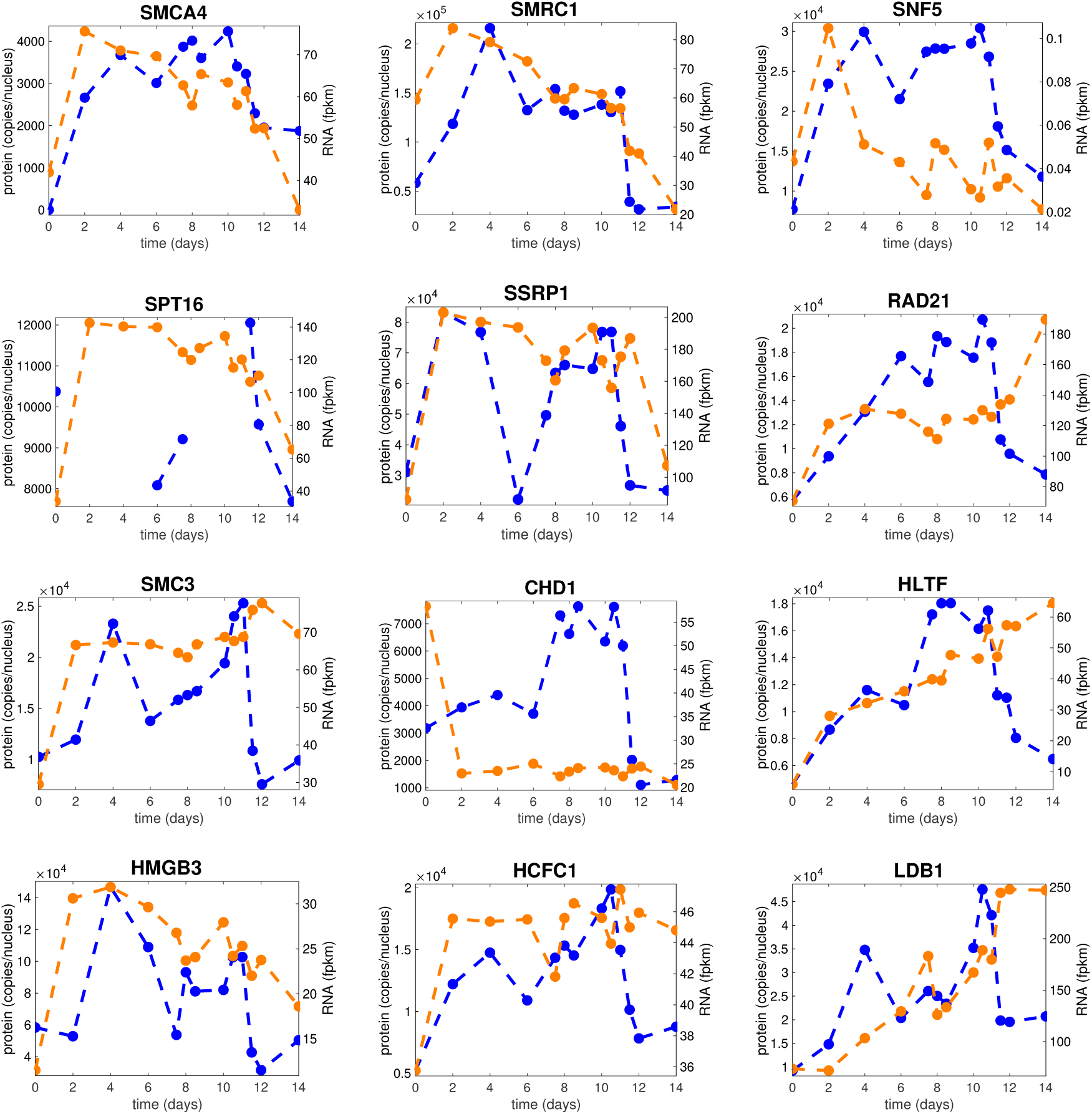
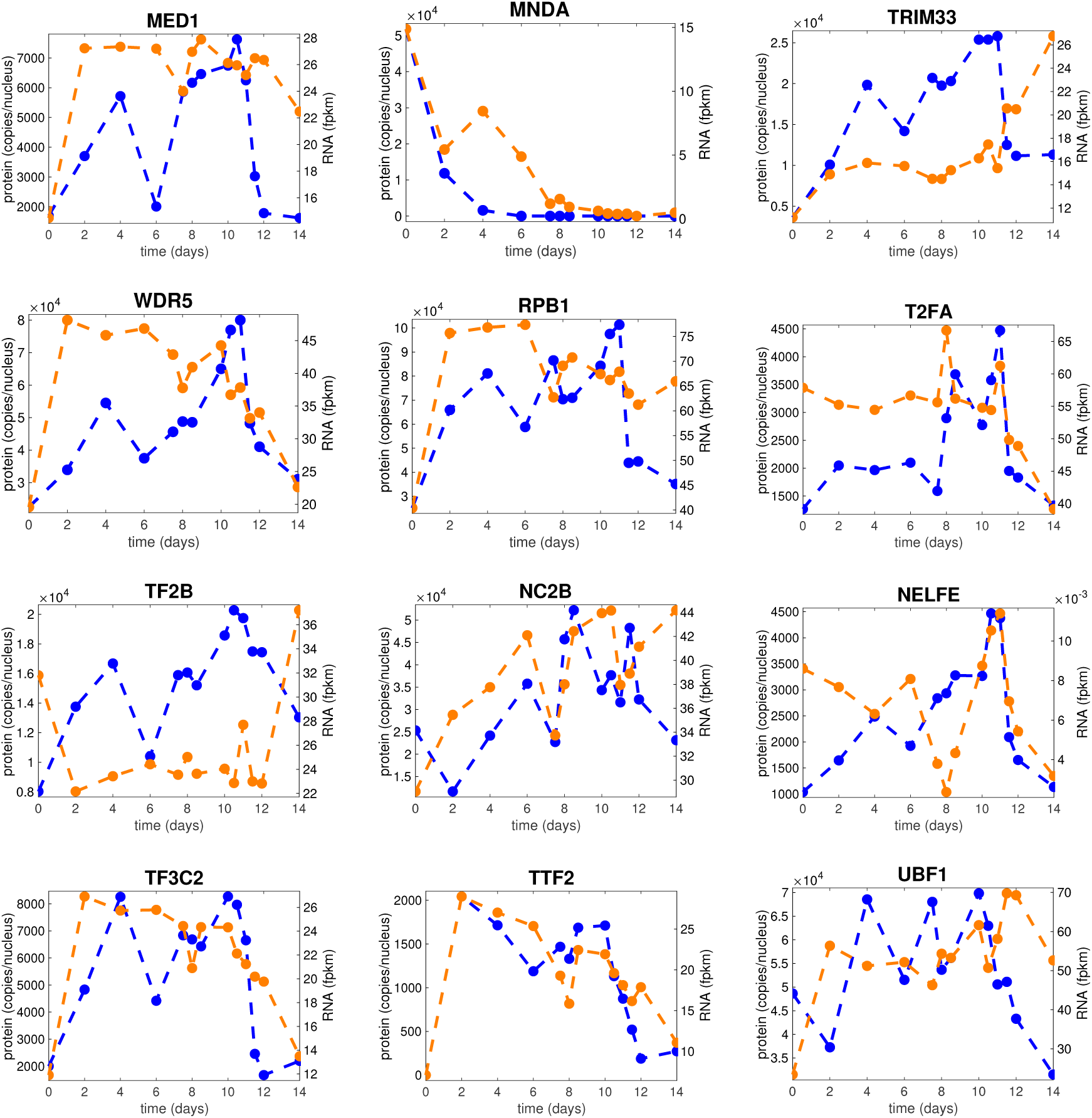
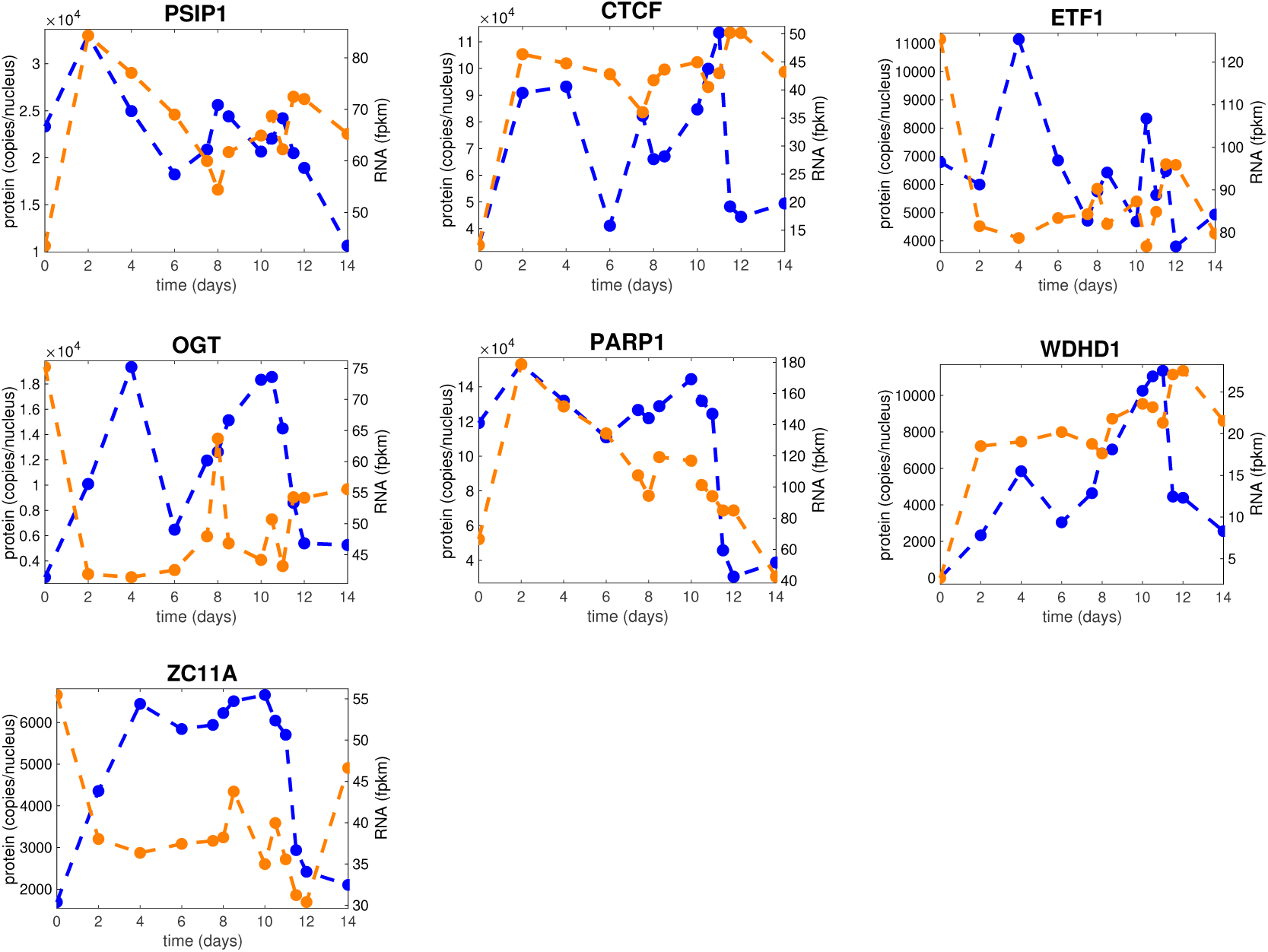
(Related to Figure 2) Comparison between mRNA and Protein Abundances during Hemato/Erythropoiesis. Protein (blue) and mRNA (orange) abundances for the indicated genes during differentiation.

**Figure S5.**
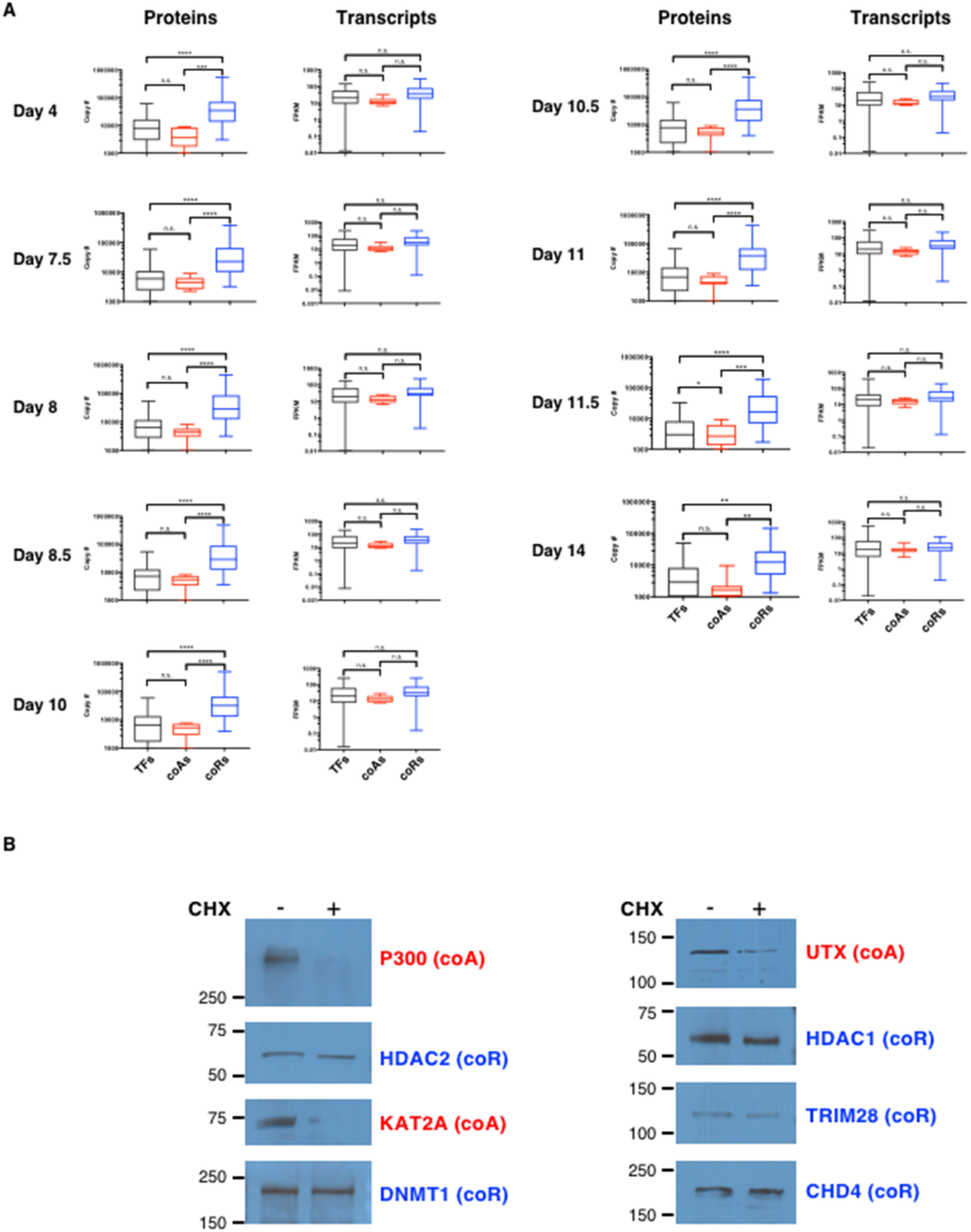
(Related to Figure 5) Co-activators are Less Stable than Co-Repressors in the Nucleus. (A) Left panels: Box plots depicting protein abundances (in copy numbers) of TFs (black), coAs (red) and coRs (blue) at the indicated days. Right panels: Box plots depicting the mRNA abundances (in FPKM) of TFs, coAs and coRs at the indicated days. Two-tailed t-test: n.s. (non-significant), * p < 0.05. **p < 0.01. *** p < 0.001. **** p < 0.0001. For a list of TF, coA and coR, see <u>Table S4</u>. (B) Western blot analyses of nuclear extracts from erythroid cells (day 8) treated with cycloheximide (CHX) to inhibit translation or a vehicle control. Molecular masses (in kDa) are indicated on the left.

## Supplemental Movie

**Movie S1.** (<u>Related to Figure 3</u>) Dynamic gene regulatory network model of erythroid commitment

### Experimental Procedures

#### Antibodies

##### Antibodies used for FACS

CD34 (CD34-PE, clone 581, BD Biosciences, cat# 555822, RRID:AB_396151), CD36 (CD36-PE, clone CB38, BD Biosciences, cat# 555455, RRID:AB_395848), CD71 (CD71-FITC, clone YDJ1.2.2, Beckman Coulter, cat# IM0483U, RRID:AB_2756301), GPA (CD235a-PE, clone GA-R2, BD Biosciences, cat# 555570, RRID:AB_395949).

##### Antibodies used for Western blot

P300 (Clone RW128, Millipore, cat# 05-257, RRID:AB_11213111), DNMT1 (Polyclonal, Abcam cat# ab87656, RRID:AB_2041078), HDAC1 (Polyclonal, Abcam cat# ab7028, RRID:AB_305705), HDAC2 (Clone Y461, Abcam cat# ab32117, RRID:AB_732777), CHD4 (Polyclonal, Abcam cat# ab72418, RRID_AB_1268107), KAT2A (L. Tora; IGBMC; Polyclonal, RRID:AB_2616158), TRIM28 (Polyclonal, LifeSpan Biosciences, cat# LS-C287177, RRID:AB_2811026), UTX (F.J. Dilworth; OHRI; Polyclonal, RRID:AB_2811027).

#### Primers for qRT-PCR

GATA2 For: 5’-CTC CCA CCT TTT CGG CTT C-3’

GATA2 Rev: 5’-CGT GGT GCT AGG GTC AGG A-3’

RUNX1 For: 5’-CCA ATA CCT GGG ATC CAT TGC-3’

RUNX1 Rev: 5’-CTG GCA CGT CCA GGT GAA A-3’

TAL1 For: 5’-CGC CTG GCC ATG AAG TAT ATC-3’

TAL1 Rev: 5’-AGG GTC CTT GCC AGT CTT-3’

FOXO3 For: 5’-CAG CCT GTC ACC TTC AGT

AAG-3’ FOXO3 Rev: 5’-TTT CAG TCA GCC CAT CAT TCA-3’

GATA1 For: 5’-AGA CTT TGA AGA CAG AGC GGC TGA-3’

GATA1 Rev: 5’-TTC CAC GAA GCT TGG GAG AGG AAT-3’

KLF1 For: 5’-GCG TTC CCA AAG ATC CAC CCA AAT-3’

KLF1 Rev: GGG TTT GCA CGA CAG TTT GGA CAT-3’

PU.1/SPI1 For: 5’-AGA AGA AGA TCC GCC TGT ACC A-3’

PU.1/SPI1 Rev: 5’-CCA CCA GAT GCT GTC CTT CAT-3’

GFI1b For: 5’-CGA CTC ACC CCC ATT CTA CAA-3’

GFI1b Rev: 5’-CGG TAG CTG TGG CCA TAG GT-3’

FLI1 For: 5’-CCA CCA ACG AGA GGA GAG TCA-3’

FLI1 Rev: 5’-CCA GCC ATT GCC TCA CAT G-3’

KLF3 For: 5’-CGA ACC ACA GAG GAC AGA TTA TT-3’

KLF3 Rev: 5’-GAC CGA AGG GTG ATT CTC TTG-3’

#### Isolation of hematopoietic stem/progenitor cells from human umbilical cord blood

Umbilical cord blood was obtained from Canadian Blood Services “Cord Blood for Research program” (CBR-2014-001). CD34^+^ hematopoietic stem/progenitor cells were isolated as previously described (Palii et al., 2011a) with the following modifications. CD34^+^ cells were pre-enriched from fresh cord blood by negative selection using the RosetteSep Human Cord Blood CD34 Pre-Enrichment Cocktail (STEMCELL Technologies cat#15631), followed by Ficoll density gradient and CD34 positive selection using the EasySep Human CD34 Positive Selection Kit (STEMCELL Technologies cat#18096) according to the manufacturer’s instructions. Purified cells were analyzed by FACS for CD34 expression using the PE Mouse Anti-Human CD34 antibody (BD Pharmingen, cat# 555822) and either cryopreserved in 10% DMSO or cultured directly as described below. All procedures were approved by the Ottawa Health Science Network Research Ethics Board (2007804-01H)

#### Human erythropoiesis ex vivo culture and cell harvest

Two biological replicates of the time-series were performed. CD34^+^ cells (63×10^6^ cells for replicate 1 and 45×10^6^ cells for replicate 2) were differentiated towards the erythroid lineage using a 4-step protocol (Giarratana et al., 2005; Palii et al., 2011a). The first step (day 0 to day 11) consists of growing CD34^+^ cells in serum-free IMDM medium supplemented with 1% penicillin/streptomycin, 4×10^-3^ M L-glutamine, 40 ug/ml inositol, 10 ug/ml folic acid, 1.6×10^-4^ M monothioglycerol, 90 ng/ml ferrous nitrate, 900 ng/ml ferrous sulfate, 20% albumin-insulin-transferrin (BIT), also containing the following cytokines: 10^-6^ M hydrocortisone (HC), 100 ng/ml stem cell factor (SCF), 5 ng/ml interleukin 3 (IL-3) and 3 IU/ml erythropoietin (EPO) for 8 days followed by 3 days in supplemented IMDM medium containing only SCF and EPO. For the second step (day 12 to day 14), cells were co-cultured on a layer of stromal MS-5 cells in the supplemented IMDM medium containing only EPO. For the third step (day 15 to day 18), cells were co-cultured on a layer of MS-5 cells in the supplemented IMDM medium with no cytokines. For the fourth step (day 19 to day 24), cells were co-cultured on a layer of MS-5 cells in the supplemented IMDM medium in the presence of 10% fetal bovine serum. Every second day, cells were counted and monitored for viability (trypan blue exclusion of dead cells), cell surface expression of CD34 (CD34-PE, BD Pharmingen, cat# 555822), CD36 (CD36-PE, BD Pharmingen, cat# 555455), CD71 (CD71-FITC, Beckman Coulter, cat# IM0483U), GPA (CD235a-PE, BD Pharmingen, cat# 555570) and LDS751 (Molecular Probes, cat# L7585) by FACS and hemoglobin production (benzidine staining). Cells were harvested at the indicated intervals during the course of differentiation and cryopreserved in their respective culture media supplemented with 10% DMSO.

#### Giemsa staining

Cells were harvested, cytospun and fixed in methanol for 5 min prior to staining with 1/20 diluted Giemsa solution (SIGMA cat# GS500) for 15 min, followed by 3 washes of 5 min each in deionized water, according to the manufacturer’s instructions.

#### RNA extraction and high throughput sequencing

For each biological replicate, total RNA was isolated on days 0, 2, 4, 6, 7.5, 8, 8.5, 10, 10.5, 11, 11.5, 12, 14 and 16 of erythroid differentiation using the RNeasy mini extraction kit (Qiagen, cat# 74104), including a DNase I digestion step. After RNA extraction, quality control was performed with RNA 6000 Nano kit (Agilent cat# 5067-1511). Libraries were prepared using a TruSeq mRNA enrich stranded RNA library kit (Illumina) with two biological replicates per library and paired-end sequencing was performed on an Illumina HiSeq 2000.

RNA seq data have been deposited in the GEO database.

#### RNAseq analysis

For each biological replicate, fastq files were aligned to the human reference genome hg38 with RefSeq annotations using Hisat2 (Kim et al., 2015). The resulting .sam files were transformed into .bam files with SAMtools (Li et al., 2009). The reads were counted using the featureCounts (Liao et al., 2014) function from the R package Rsubread, specifying the same .gtf file used to build the Hisat2 index and default parameters. Differentially expressed genes were identified with the R package DESeq2 (Love et al., 2014). Genes with adjusted p-value below 0.05 were considered statistically significant. For visualization the data was normalized to FPKMs. Principal Component Analysis was done with the R function prcomp using the log(1+FPKM) transformed data.

#### Nuclear protein extraction and relative quantification using iTRAQ

Fifteen million cryopreserved cells at days 0, 2, 4, 6, 8, 10, 12 and 14 of erythroid differentiation were thawed and resuspended in serum-free IMDM medium supplemented with 1% penicillin/streptomycin, 4×10^-3^ M L-glutamine, 40 ug/ml inositol, 10 ug/ml folic acid, 1.6×10^-4^ M monothioglycerol, 90 ng/ml ferrous nitrate, 900 ng/ml ferrous sulfate, 20% albumin-insulin-transferrin (BIT). After thawing, the cells were washed twice with ice-cold PBS, resuspended in ice-cold Swelling Buffer (10mM HEPES pH7.9, 1.5mM MgCl_2_, 10mM KCl, 0.1% (v/v) NP-40, protease inhibitor cocktail) and incubated on ice for 30 min. During incubation, cells were vortexed every 5 min to allow cell lysis. Nuclei were then pelleted by centrifugation at 1,500 rpm (4°C) for 5 min, washed twice with ice-cold PBS and resuspended in RIPA Buffer (50mM HEPES pH7.9, 1mM MgCl_2_, 150mM NaCl, 0.5% (w/v) Na deoxycholate, 1% (v/v) NP40, 0.1% SDS) containing 50 ng/µl Benzonase (Millipore, cat# 70746) and protease inhibitor cocktail at room temperature (RT). Samples were vortexed for 5 min at RT and incubated for 20 min at 37°C on a Thermomixer (14,000 rpm) followed by 5 min vortexing at RT. Nuclear extracts were recovered by centrifugation at 14,000 rpm for 15 min, snap frozen in liquid nitrogen and stored at -80°C. Nuclear protein extracts were prepared from cells on days 0, 2, 4, 6, 8, 10, 12, and 14 of erythroid differentiation. Extracted protein concentrations were measured using the bicinchoninic acid assay (BCA; Thermo Scientific). Equal protein amounts were reduced with 5mM dithiothreitol, alkylated with 10mM iodoacetamide, and precipitated with chilled 100% acetone to remove detergent. Precipitated proteins were resuspended in iTRAQ Dissolution Buffer (AB Sciex) and digested into solution with Lys-C for 3 h (1:200 w:w, 37C; Thermo Scientific) followed by Trypsin overnight (1:50 w:w; 37C; Thermo Scientific). Volatile liquids were removed by evaporation, and peptides were resuspended in iTRAQ Dissolution Buffer and labeled with 8-plex iTRAQ reagents (AB Sciex) for 2 h at room temperature with constant agitation. The labeling reaction was quenched with 1M Tris, pH 8.0, and samples were combined and volatile liquid was evaporated. Labeled peptides were resuspended in 1% acetonitrile, acidified with formic acid, and purified using C18 reversed-phase chromatography (1cc 100mg cartridges; Waters). Purified peptides were separated into 24 fractions using isoelectric focusing off-gel electrophoresis (Agilent), and ampholytes removed by tC18 reversed-phase chromatography (100mg 96-well plate; Waters) followed by mixed cation exchange chromatography (30um uElution 96-well plate; Waters). Purified peptides were separated by online nanoscale HPLC (EASY-nLC1000; Thermo Scientific) with a C18 reversed-phase Picochip nanospray column pre-packed 10.5cm with ReproSil-Pur C18-AQ 3um 120A (New Objective) over an increasing 90 min gradient of 5-35% Buffer B (100% acetonitrile, 0.1% formic acid) at a flow rate of 300nl/min. Eluted peptides were analyzed with a Q Exactive HF Hybrid Quadrupole-Orbitrap mass spectrometer (Thermo Scientific) operated in data dependent mode, with the Top15 most intense peptides per MS1 survey scan selected for MS2 fragmentation by higher energy collisional dissociation (HCD). MS1 scans were performed in the Orbitrap at a resolution of 60,000 at m/z 400, with an automatic gain control (AGC) target of 3e6 ions and a maximum injection time of 20ms. MS2 scans were analyzed in the Orbitrap at a resolution of 15,000, with an AGC target of 1e5 and a maximum injection time of 45ms. Due to the iTRAQ label, a fixed first mass of 100 m/z was used for MS2 scans, along with a normalized collision energy of 30 and an isolation window of 1.2 m/z. Peptide match was not used, and dynamic exclusion was set to 30 s (+/-15 ppm). Raw output data files were searched against the Uniprot human protein database (12-2015 release) using X!Tandem and Comet (Craig and Beavis, 2004; Eng et al., 2013) to identify peptide sequences. A reverse sequence database was appended to assist in determining error. Resulting data were combined using iProphet, and probabilities for correct identification were determined by Peptide prophet, iProphet, and Protein prophet (Keller et al., 2002; Nesvizhskii et al., 2003; Shteynberg et al., 2011). iTRAQ quantification was performed using Libra (Pedrioli et al., 2006).

iTRAQ data has been deposited in ProteomeXchange via MassIVE.

#### iTRAQ data analysis

iTRAQ data was clustered using k-means clustering with the R function kmeans, with centers=8 and default parameters. Gene Ontology (GO) analysis of the different clusters was done using Metascape (Zhou et al., 2019) with the option Express Analysis.

#### Construction of an Erythroid protein database

Day 8 nuclear protein extracts (86 µg) were reduced with 5mM dithiothreitol, alkylated with 25mM iodoacetamide, and digested with Lys-C for 3 h (1:200 w:w, 37C; Thermo Scientific) followed by Trypsin overnight (1:25 w:w, 37C; Thermo Scientific). Peptides were acidified with formic acid, purified using C18 reversed-phase chromatography (1cc 100mg cartridges; Waters), and separated into 24 fractions using isoelectric focusing off-gel electrophoresis (Agilent). Ampholytes were removed by tC18 reversed-phase chromatography (100mg 96-well plate; Waters) followed by mixed cation exchange chromatography (30um uElution 96-well plate; Waters). Purified peptides were separated by online nanoscale HPLC (EASY-nLC II; Proxeon) with a C18 reversed-phase column packed 25cm (Magic C18 AQ 5um 100A) over an increasing 60 min gradient of 5-35% Buffer B (100% acetonitrile, 0.1% formic acid) at a flow rate of 300nl/min. Eluted peptides were analyzed with an Orbitrap Elite mass spectrometer (Thermo Scientific) operated in data dependent mode, with the Top20 most intense peptides per MS1 survey scan selected for MS2 fragmentation by rapid collision-induced dissociation (rCID) (Michalski et al., 2012). MS1 survey scans were performed in the Orbitrap at a resolution of 240,000 at m/z 400 with charge state rejection enabled, while rCID MS2 was performed in the dual linear ion trap with a minimum signal of 1000. Dynamic exclusion was set to 15 s (+/-10 ppm). Raw output data files were searched against the Uniprot human database (03-2015 release) using X!Tandem and Comet (Craig and Beavis, 2004; Eng et al., 2013) to identify peptide sequences. A reverse sequence database was appended to assist in determining error. Resulting data were combined using iProphet, and probabilities for correct identification were determined by Peptide prophet, iProphet, and Protein prophet (Keller et al., 2002; Nesvizhskii et al., 2003; Shteynberg et al., 2011). Xpress was used to determine the summed peak area for each peptide (Han et al., 2001).

#### Construction of the Erythroid SRM TF Atlas

We created a list of 168 TFs with known and/or expected roles in transcriptional regulation during erythropoiesis. Proteotypic, fully tryptic peptides with the highest summed MS1 peak areas were selected from the erythroid protein database for all proteins of interest. In addition, selected peptides were a minimum of 7 amino acids in length and lacked features that may be incompatible with SRM analysis, namely ragged ends or missed cleavages. Where possible, Cys-containing peptides, peptides containing potentially modified residues (*e.g.*, Met, Ser, Thr, Tyr, N-terminal Gln, Asn-Gly, Gln-Gly) and sequences that could affect trypsin digestion efficiency or peptide stability were also avoided. For proteins not found in our database, or to supplement our identified peptides, the SRM Atlas (Kusebauch et al., 2016) was used with a preference for peptides validated on an Agilent triple quadrupole mass spectrometer. Transition selection and collision energy values were also imported from the SRM Atlas. SIL peptides (Lys-[13C6, 15N2] or Arg-[13C6, 15N4]) were synthesized for all proteins (716 peptides; PEPotec Grade 1; Thermo Scientific). These SIL peptides were used as standards for confident peptide identification during SRM and to optimize SRM assays. Assays were refined to include those transitions which displayed the highest interference-free signal intensities in solvent and when added to erythroid cell extracts. This process resulted in 3-4 transitions per peptide (and 1-4 peptides per protein). The linear range of quantification within the erythroid cell extracts was determined for the SIL peptides used for quantification. The concentration of SIL peptides used in the final time course measurements was matched to that detected in day 8-12 erythroid nuclear extracts. The final erythroid SRM TF Atlas consists of 150 proteins, 411 heavy and light peptide pairs and 1377 heavy and light transition pairs.

#### Nuclear protein extraction and sample preparation for SRM analyses

Fifteen million cryopreserved cells at days 0, 2, 4, 6, 7.5, 8, 8.5, 10, 10.5, 11, 11.5, 12, and 14 of erythroid differentiation were thawed and washed using IMDM supplemented medium. Cells were then washed in ice-cold PBS buffer, resuspended in ice-cold Swelling Buffer (10 mM HEPES pH7.9; 1.5 mM MgCl_2_; 10 mM KCl; 0.1% (v/v) NP40; protease inhibitor cocktail) and incubated on ice for 30 min. During incubation, cells were vortexed every 5 min to allow cell lysis. Nuclei were then pelleted by centrifugation for 5 min at 1,500 rpm (4°C) and resuspended in 1 vol. of 37°C pre-heated Extraction Buffer 1 (50 mM HEPES pH7.9; 1 mM MgCl_2_; 150 mM NaCl; 0.5% Na deoxycholate; 50 ng/µl Benzonase Millipore, cat# 70746; protease inhibitor cocktail) prior to incubation at 37°C on a Thermomixer (14,000 rpm) for 15 min. Proteins were extracted first by 6 passages through a 27 ½ gauge needle prior to addition of 1 vol. of 37°C pre-heated Extraction Buffer 2 (50 mM HEPES pH7.9; 150 mM NaCl; 9.5% Na deoxycholate; 1 mM EDTA; protease inhibitor cocktail). The mixture was heated at 70°C for 5 min and proteins were extracted further by 6 passages through a 27 ½ gauge needle prior to incubation at 40°C on a Thermomixer (14,000 rpm) for 15 min. Nuclear extracts were recovered by centrifugation at 13,000 rpm for 15 min and snap frozen. Extracted protein concentrations were measured using the BCA assay (Thermo Scientific). Equal protein amounts were denatured by boiling at 100°C for 4 min, cooled and reduced with 5mM dithiothreitol, alkylated with 25mM iodoacetamide, and digested with Lys-C for 3 h (1:200 w:w, 37C; Thermo Scientific). Samples were diluted to reduce sodium deoxycholate concentration to below 1%, and digested with Trypsin overnight (1:25 w:w, 37C; Thermo Scientific). Concentration-matched isotopically heavy peptide standards were added to erythroid peptide samples after overnight digest. To remove sodium deoxycholate and prepare peptides for purification, samples were acidified with an equal volume of cold 1% trifluoroacetic acid. Acidified supernatants were subsequently purified by mixed cation exchange chromatography (30um uElution 96-well plate; Waters).

#### SRM analysis

SRM was performed on an Agilent 6490 triple quadrupole mass spectrometer, equipped with a chip cube interface. Peptides were separated by online HPLC (1260 Infinity; Agilent) with a reversed-phase microfluidics HPLC chip (160nL trap; Agilent) over an increasing 60min gradient of 3-25% acetonitrile. Optimized transitions were acquired in dynamic MRM mode, with a 5 min retention time window, using MS1 wide and MS2 unit resolutions. Collision cell accelerator voltage was set to 5V, and cycle times were set to yield a minimum dwell time of 12ms. Raw data was processed using Skyline (MacLean et al., 2010) and peaks were manually verified. Light-to-heavy ratios were calculated from peak area values. Only measurements determined to be within the linear range of quantification for the mass spectrometer were used for subsequent analyses. SRM data has been deposited into The Peptide Atlas SRM Experiment Library (PASSEL).

#### Absolute quantification of transcription factors by SRM

Absolute quantification was achieved in two ways: The first approach (71 proteins) is based on stable isotope dilution (SID) in which peptide abundance is determined by comparison of selected transition peak areas for each peptide to those of its corresponding SIL peptide. The second approach (34 proteins) is a label free method in which SID is used to determine the concentration of a set of 17 “anchor” proteins, and standard curves based on transition peak areas from the two highest intensity peptides per anchor protein and their concentrations are then used to estimate the concentrations of the target proteins (Ludwig et al., 2012). For SID, isotopically light internal standard (IS) peptides (Gerber et al., 2003) were used to quantify our SIL standard peptides. This approach enabled a significant time, cost and resource savings versus using isotopically heavy IS peptides for assay development and direct quantification in the biological matrix. Peptides that showed the highest interference-free intensity were selected for commercial synthesis (AQUA; Thermo Scientific). IS peptides were >97% pure and were quantified by amino acid analysis. For SRM quantification, the peak areas for the two most intense, interference-free transitions were summed together. Additional transitions were used to confirm correct peak identification. Dose-response curves were generated for each IS peptide at abundances ranging from 10 amol to 500 fmol. Quantification of the SIL peptides was achieved by measuring transition intensities from serial dilutions of the peptides and then plotting the corrected peak areas onto the corresponding IS dose-response curve. The light-to-heavy ratios calculated for the erythroid differentiation time course were used to convert these SIL standard abundance measurements into endogenous protein abundances. Peptide-based protein abundances were averaged in cases where more than 1 peptide/protein was quantified.

For label free absolute quantification, the following criteria were used to select “anchor” proteins for the standard curves: (i) IS dose-response regression line was linear between 0.1-500 fmol with an R2>0.98; or (ii) IS dose-response regression line was linear between 1-500 fmol with an R2>0.98 and a slope>0.8; or (iii) IS dose-response regression line was linear between 1-500 fmol with an R2>0.95 and the peptide was included based on (i) or (ii) in another replicate. Peak areas were calculated by summing the two most intense interference-free transitions per peptide, and averaging these values for the two highest intensity peptides per protein. Linear regression was used to fit a standard curve to the above values, and to estimate the unknown concentrations of endogenous proteins using their measured transition peak area values.

SRM measurements obtained as described above produced biological duplicate values in units of fmol/µg of protein for each protein at each time point. The two replicates were combined by the following steps, to produce a single abundance value for each gene at each time point. First, if both replicates had a valid measurement, the two were averaged. If only one replicate had a valid measurement, it alone was taken as the representative value. If neither replicate had a valid measurement, the value was marked as missing. Second, missing values were filled in by linear interpolation where possible. For each missing value (say on day y), we sought the latest non-missing value prior (say on day x) and the earliest non-missing value later (say on day z). If such days could be found, then the abundance at day x was filled in with (z-y)/(z-x) times the abundance at day x, plus (y-x)/(z-x) times the abundance at day z. This interpolation rule was not applied to genes ETO2, MLL1 and SPT16, as visual inspection suggested there were too many missing values and/or not clear enough of a trend in abundance for interpolation to be meaningful. This left only one gene, BACH1, with a missing value at our final time point, which we filled in as zero consistent with BACH1’s general trend towards decreasing expression. See <u>Table S4</u>. Some analyses, such as the GRN modeling, used these numbers. For other analyses, we wanted to express protein abundance in units of protein copy number per nucleus. To obtain these numbers, we multiplied by 4420.25799, which comes from the following considerations: (1) There are 6.022140857e+8 molecules per fmol. (2) Across all samples, we extracted an average of 7.3426 µg protein per million nuclei.

#### Protein and RNA correlation analyses

Correlation heatmap values were computed with the cor R function using Pearson correlation, computing the correlation over all genes on FPKM normalized data. The heatmaps were plotted using a custom Matlab script.

#### iTRAQ versus SRM correlation

Correlation analysis across time points by computing the Pearson correlation of iTRAQ relative protein abundance and SRM absolute protein abundance (copy number) for each gene. The histogram of the correlation was done with the R function *hist* with breaks=20.

#### Protein versus mRNA correlation analyses

Correlation analysis across genes was performed by computing the Spearman correlation of the mRNA abundance (FPKM) and protein abundance (copy number) of all genes at each time point. Correlation analysis over time was performed by computing the Pearson correlation of the mRNA abundance (FPKM) over time with the respective protein abundance (copy number) for each gene over time.

Further mRNA and protein correlation analyses can be performed on our Human Erythropoiesis TFs website: https://trena.systemsbiology.net/app/srm_rna_combined_v2

#### Gene Regulatory Network modeling

For a core set of factors (ELF1, ERG, FLI1, GATA1, GATA2, GFI1B, KLF1, NFE2, RUNX1, TAL1, SPI1), possible regulatory links were obtained from the literature (Dore and Crispino, 2011; Gottgens, 2015; Sive and Gottgens, 2014). For three genes (E2F2, HXB4, KLF3), literature evidence was insufficient to suggest regulatory links. For each of these three genes, we computed correlations between the RNA expression of the gene and the protein expression of all other genes in the network. The three other genes with highest correlation were posited to be possible activators, and the gene with most negative correlation was posited to be a possible repressor. This network of possible activators and repressors was further refined by fitting ordinary differential equation models to the quantitative expression data.

Given a gene X, let x denote our modeled (or predicted) RNA expression of gene X, in units of FPKM. Let A_i_(t) denote the observed protein expression of the i^th^ activator of gene X, and R_j_(t) the observed protein expression of the j^th^ repressor of gene X, both in units of fmol/µg. We model the RNA transcription of X as

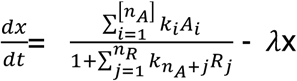

where k_i_ and λ are constants to be determined, *n_A_* is the number of activators considered for X and *n_R_* is the number of repressors considered for X. Given an initial condition, x(0), values for the A_i_ and R_i_ as a function of time, and values for the regulatory and decay parameters, the above ordinary differential equation (ODE) can be solved numerically over any finite time interval, producing predicted values x(t, k, λ). We used a Runge-Kutta 7(8) method to calculate such trajectories. The values for for A_i_ and R_j_ as a function of time are obtained by interpolation from the observed SRM (protein) values, using a cubic spline smoothing from the GSL libraries (Galassi).

Given also observed RNA expression values for gene X, O(t_i_) at observation times t_1_, t_2_, …, t_m_, agreement between predicted and observed expression values can be quantified using the objective function

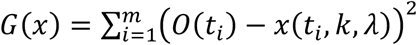

To find regulatory and decay parameters, we sought to minimize the objective function, subject to the constraints that all parameters need to be positive and decay rate has to be larger than 16.6 day^-1^.

The minimum of the objective function was computed using a Simplex method from the GSL libraries. To explore the parameter space we choose random initial parameters between 0 and 500 for the k’s, and between 16.6 day^-1^ and 500 day^-1^ for the λ with stop conditions either error below 1e-04 or a maximum of 2000 iterations. With a total of 128 random initial conditions. Out of the 128 iterations, the parameter set that gives the smallest values in the objective function were selected.

#### Gene Regulatory Network display

For activators, the transparency score of the link representing A_i_ activating X at time t is

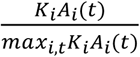

Links with transparency score below 0.1 at a certain time are not displayed. Links with transparency scores between 0.1 and 1 are increasingly opaque, and links with scores greater or equal to 1 are fully opaque. For repressors, the transparency score of the link representing R_i_ repressing X is log_2_(1+K_i_R_i_). The thickness of a link from a regulator (activator A_i_ or repressor R_i_) to gene X is proportional to log(1+K_i_).

The network can be further explored in BioTapestry format (Paquette et al., 2016) at the following link: http://grns.biotapestry.org/HumanErythropoiesisGRN/

#### Intersection of GRN putative links with published ChIP-seq data

Published ChIP-seq data for ERG (GSM1097879), FLI1 (GSM1097880), GATA1 (GSM1278240, GSM1816081, GSM970258, GSM1816080, GSM970257), GATA2 (GSM1097883, GSM1816093, GSM722396), GFI1B (GSM1278242), KLF1 (GSM2575041-GSM2575049), RUNX1 (GSM1816092, GSM1097884, GSM1816091) and TAL1 (GSM1278241, GSM1816084, GSM1097881, GSM1427077, GSM1816082, GSM1816083) corresponding to CD34^+^ HSPC or ProEB stages (Beck et al., 2013; Huang et al., 2016; Norton et al., 2017; Pinello et al., 2014; Trompouki et al., 2011; Xu et al., 2012; Xu et al., 2015) were downloaded from NCBI GEO and analyzed as follow. Raw fastq data was trimmed for low quality bases using trimmomatic (version 0.38) (Bolger et al., 2014), and mapped to the human genome (hg38) using bowtie-2 (Langmead and Salzberg, 2012). Aligned reads were filtered for multiple mapping using a mapping quality filter (Q20) and the filtered alignments were sorted (using samtools version 1.9 (Li et al., 2009)) and used for calling peaks. Peaks were called using MACS2 (version 2.1.2) (Zhang et al., 2008), with a peak enrichment cut-off filter of P = 0.05. The peaks were filtered against ENCODE blacklist (Amemiya et al., 2019) for human genome using bedtools v2.27.1. For each sample, an appropriate Input sample from the same stage of development was used as a control. Genome coverage files were created using deepTools2 suite (bamCoverage) (Ramirez et al., 2016). The identified ChIP-seq peaks were overlapped with GeneHancer (v4.4) (Fishilevich et al., 2017) elements (promoters and enhancers) using bedtools. This filtered list was then intersected with putative links from the GRN model at day0 (HSPCs) or day 10 (ProEB) to identify possible direct interactions between TFs and target genes (TGs). Based on this data, we classified TF–TG pairs as either “interacting” or “not interacting” and calculated the percentage of regulatory links that could be explained by direct TF binding.

#### Lentivirus preparation and infection

Lentiviral particles expressing shRNA sequences against GATA2 (5’- CCAGACGAGGTGGACGTCTTCTTCAATCA-3’), GATA1 (5’- GATCCCCGAAGCGCCTGATTGTCAGTTTCAAGAGAACTGACAATCAGGCGCTTCTTTTTGGAAA-3’), TAL1 (5’- CTTACTCTAGGAGGCGGAC-3’), and KLF1 (5’- CCGGACACACAGGATGACTTCCTCAAGTG -3’) were prepared as previously described (Palii et al., 2011a). Specifically, 293T cells were transfected with the pMD2.G envelope vector (Addgene #12259), the psPAX2 packaging vector (Addgene, #12260) and one of the following shRNA expression lentiviral vectors: GATA2 shRNA Lentivector Target a (Abm #i008537a), pLVUTHshGATA1-tTR-KRAB (Addgene #11650), pBLOCK-it6-DEST (sh Tal1) (Palii et al., 2011b) or KLF1 shRNA Lentivector Target a (Abm #i011644a), using calcium phosphate precipitation. Lentiviral particles were harvested, concentrated by ultracentrifugation (50,000 g for 2h) and used to infect cells at the day 8 time-point with a MOI of 20. Lentiviral infection was repeated 24h later in the same conditions. Cells were harvested 24h after the last infection and used for RNA extraction.

#### ATACseq, HINT-ATAC and estimation of the number of enhancers

ATAC-seq was performed as previously described (Buenrostro et al., 2013; Hay et al., 2016) for cells at days 8, 10 and 12. Briefly, 75000 cells per technical replicate per sample per timepoint were lysed in cold lysis buffer, nuclear pellets were obtained after 10 min centrifugation at 4°C at 500G and resuspended in 50ul of tagmentation mix (FC-121-1030, Illumina), then incubated for 30 min at 37°C. DNA was purified using the Qiagen MinElute columns (28004, Qiagen). Tagmented DNA was indexed with custom primers using NEB Next High-Fidelity 2x PCR Master Mix (M0541S, NEB). And purified with Qiagen PCR Cleanup Kit (28104, Qiagen). Samples were multiplexed sequenced on a next generation sequencing platform using the NextSeq^®^ 500/550 High Output Kit v2 (75 cycles; FC-404-2005, Illumina) using paired-end reads. For data analysis, the fastq data was mapped onto the human genome (hg38) using bowtie1.0 (Langmead et al., 2009) with the following parameters: *--chunkmb 256 –phred-quals 33 –m 2 –best –strata –maxins 400*. The mapped bam files were used to call narrow peaks using MACS2 with the version 2.1.2 docker image of macs https://hub.docker.com/r/fooliu/macs2

In preparation for running HINT-ATAC, all peaks which mapped to non-canonical chromosomes (chr1-22, X, Y, M) were eliminated (“cleaned”) and a bam index was built for each bam file using samtools version 1.8. HINT was then run from the regulatory genomics toolkit, version 0.12.3 (http://www.regulatory-genomics.org/hint/introduction/) on the indexed bam file, and the cleaned peaks file rgt-hint footprinting --atac-seq --organism=hg38 $(BAM) $(PEAKS) Putative enhancer and promoter regions from GeneHancer 4.1.1. were identified for all tissues, and the Bioconductor GenomicRanges package was used to calculate the intersection of macs2 narrowpeaks and hint-called ATAC-seq footprints with the sets of enhancers.

ATAC seq data have been deposited in the GEO database.

#### Cycloheximide treatment and nuclear protein extraction

Erythroid cells at day 8 were treated with 1μg/mL of cycloheximide or DMSO vehicle control for 3h. Cells were washed twice with ice-cold PBS, resuspended in ice-cold Swelling Buffer (10mM HEPES pH7.9, 1.5mM MgCl_2_, 10mM KCl, 0.1% (v/v) NP-40, protease inhibitor cocktail) and incubated on ice for 30 min. During incubation, cells were vortexed every 5 min to allow cell lysis. Nuclei were then pelleted by centrifugation at 1,500 rpm (4°C) for 5 min, washed twice with ice-cold PBS and resuspended in RIPA Buffer (50mM HEPES pH7.9, 1mM MgCl_2_, 150mM NaCl, 0.5% (w/v) Na deoxycholate, 1% (v/v) NP40, 0.1% SDS) containing 50 ng/µl Benzonase (Millipore, cat# 70746) and protease inhibitor cocktail at room temperature (RT). Samples were vortexed for 5 min at RT and incubated for 20 min at 37°C on a Thermomixer (14,000 rpm) followed by 5 min vortexing at RT. Nuclear extracts were recovered by centrifugation at 14,000 rpm for 15 min, snap frozen in liquid nitrogen and stored at -80°C before Western blot analysis.

